# Taxonomically different symbiotic communities of sympatric Arctic sponge species show functional similarity with specialization at species level

**DOI:** 10.1101/2025.03.15.643485

**Authors:** Anastasiia Rusanova, Viktor Mamontov, Maxim Ri, Dmitry Meleshko, Anna Trofimova, Victor Fedorchuk, Margarita Ezhova, Yulia Lyupina, Alexander Finoshin, Artem Isaev, Dmitry Sutormin

## Abstract

Marine sponges harbor diverse communities of associated organisms, including eukaryotes, viruses, and bacteria. Sponge associated microbiomes contribute to the health of the host organisms by defending them against invading bacteria and providing them with essential metabolites. Here we describe microbiomes of three sympatric species of cold-water marine sponges – *Halichondria panicea*, *Halichondria sitiens*, and *Isodictya palmata* – sampled over a period of six years at the White Sea. We identified the sponges as low microbial abundance species and detected stably associated bacteria that represent new taxa of sponge symbionts within Alpha- and Gammaproteobacteria. The sponges carried unique sets of unrelated species of symbiotic bacteria illustrating varying complexity of microbiomes. On a community level, sponge associated microbiomes shared common symbiotic features; they encoded multiple eukaryotic-like proteins, biosynthetic pathways, and transporters of amino acids and vitamins essential for sponges. On a species level, however, different classes of eukaryotic-like proteins and pathways were distributed between dominant and minor symbionts indicating specialization within microbiomes. Particularly, taurine and sulfoacetate metabolism pathways were associated exclusively with dominant symbionts in all three sponge species. Our study demonstrates strong functional convergence and co-evolution of microbiomes of sympatric cold-water sponge species with a distribution of functions between community members.

Additionally, we observed dramatic shifts in compositions of sponge microbiomes coinciding with abnormally high water temperatures during the 2018 season, highlighting the vulnerability of cold-water ecosystems to global warming.

## Introduction

Sponges, phylum Porifera, exhibit extensive global distribution across a wide range of habitats and display remarkable biodiversity [1]. Sponges host complex communities of eukaryotes, prokaryotes, and viruses comprising a holobiont, wherein each participant contributes to the stability and functionality of the entire system [2, 3]. Based on diversity and abundance of sponge associated microbiomes (SAMs), sponge species are categorized as either high or low microbial abundance (HMA or LMA, respectively). HMA sponges can host more than a dozen distinct bacterial phyla with a total density reaching 10^8^–10^10^ bacteria per gram of sponge wet weight, whereas less complex microbiomes of just 2–5 phyla with a total density of 10^5^–10^6^ are typical for LMA sponges [4–6]. To date, 40-60 different bacterial phyla have been identified in various sponge species [7, 8]. Sponge-associated bacteria (SABs) are generally divided into transiently associated and stably associated groups. The latter are typically enriched in sponges compared to the surrounding water and exhibit high host specificity. Stably associated SABs are usually referred to as sponge symbionts, although functional characterization of host-bacterial interactions remains limited [9–11]. Such specificity makes the cultivation of sponge symbionts particularly challenging, as these bacteria are adapted to life within sponge tissues or cells. Consequently, metagenomics and metatranscriptomics serve as key tools for studying SAMs and predicting their functional traits [11].

Metagenomic studies showed that most bacterial symbionts from different sponge species and various geographic locations share specific genomic traits (usually called symbiotic features) that reflect adaptation to a host and contribute to the stability of a holobiont. Among these features are the biosynthetic pathways and transporters of vitamins and amino acids, which are encoded in the genomes of sponge symbionts. It is suggested that bacteria provide these compounds to hosts [3, 12, 13]. At the same time, symbionts likely obtain certain nutrients from a host. For instance, symbiotic bacterium *Candidatus* Taurinisymbion ianthellae is likely capable of utilizing taurine, potentially produced by a sponge, as a source of carbon, sulfur, and energy [14]. Another example of symbiotic features includes genes of eukaryotic-like proteins (ELPs) which are enriched in SAM metagenomes and genomes of symbiotic bacteria. ELPs are believed to mediate interactions with a host and be responsible for the recognition of symbionts and their discrimination from food-source bacteria [15–17].

Historically, most research on sponge microbiomes focused on species from tropical and temperate regions, while polar sponges remained relatively understudied, with a growing interest emerging only in recent years [18–23]. Particularly, studies highlight the vulnerability of cold-water sponges and their microbiomes to global warming, which is an increasing concern [24–27]. To study the long-term stability of polar SAMs and their functional traits, we applied metagenomics to three common sympatric sponge species (*Halichondria panicea*, *Halichondria sitiens*, and *Isodictya palmata*) from the White Sea over the sampling period of 6 years (2016-2022). Using 16S metagenomics, we identified stably-associated bacterial operational taxonomic units (OTUs) enriched in sponge microbiomes. We found fluctuations in the composition of SAMs, affected by anomalously increased water temperatures during the 2018 summer season and recovered by 2022. Using the hybrid shotgun approach and the state-of-the-art CORe contigs ITerative Expansion and ScaffoLding refining algorithm (CORITES), we obtained high-quality metagenome-assembled genomes (MAGs) of the symbiotic bacteria. Phylogeny reconstruction assigned identified symbionts to six novel species of Proteobacteria: *Ca*. Yagmuria paniceus, *Ca*. Ahtobacter symbioticus, *Ca*. Vellamobacter salmiensis, *Ca*. Vienanmeria sitiensis, *Ca*. Sampovibrio pertsovi, and *Ca*. Eurynomebacter symbioticus. Using genomic analysis, we revealed converging metabolic specializations within communities for a symbiotic lifestyle. Finally, with fluorescence *in situ* hybridization (FISH) microscopy, we verified the presence of identified symbiotic bacteria within sponge tissues and found species-specific localization of bacterial cells.

## Materials and methods

### Sample collection

For metagenomics, fragments of visually healthy *Isodictya palmata* (Ellis & Solander, 1786), *Halichondria sitiens* (Schmidt, 1870), and *Halichondria panicea* (Pallas, 1766) marine sponges were collected by SCUBA divers at 5-7 m water depth at the N. Pertsov White Sea Biological Station (WSBS MSU, 66.5527°N, 33.1033°E, Kandalaksha Bay of the White Sea, Russia) in August-September of 2016, 2018, and 2022 (**Supplementary Table S1**). Sponge species were held separately for 2 h at 5°C in 5 L of seawater sterilized by filtering through a 0.22 μm filter (Sartorius). Identification of sponge species was performed by zoologist Dr. Boris Osadchenko (Lomonosov MSU) and further confirmed by 18S rRNA gene region amplification and sequencing (**Supplementary Table S2**). As a control, 3 L of surrounding seawater at the sampling site was collected in a sterile container and was immediately processed in the WSBS laboratory. Specimens of *Halichondria panicea* were also collected from the sublittoral zone at three different sites around Dalnye Zelentsy (69.10177 N 36.06234 E) the Barents Sea, Russia) in August 2022 (**Supplementary Table S1**).

### Microbiome 16S and shotgun sequencing data analysis

Isolation of microbiomes from sponge samples and seawater, DNA extraction, and sequencing were performed as described in **Supplementary Methods**. 16S rRNA raw data analysis was performed as described earlier [28]. Briefly, trimmed forward reads were processed with DADA2 pipeline [29] and resultant amplicon sequence variants (ASVs) were clustered with MMseqs2 [30] into operative taxonomic units (OTUs). Taxonomy was assigned to OTUs using SILVA [31]. PCoA (Principal coordinates analysis), alpha-diversity, and taxonomic analyses were performed with the phyloseq package [32]. To identify SABs, OTUs meeting the following criteria were selected: (a) observed in most of the sponge samples of a particular species; (b) with a relative abundance exceeding 1% in sponge samples; (c) with a relative abundance at least 50 times higher in a sponge sample than in seawater.

The methodology for assembly, annotation, and binning of shotgun metagenomes is described in detail in **Supplementary Methods**. Briefly, Illumina or BGI reads were assembled with SPAdes [33]. Nanopore reads were assembled with Flye [34]; SPAdes was used for hybrid assembly.

Obtained assemblies were binned with MaxBin 2.0 [35], CONCOCT [36], MetaBAT 2 [37], and binny [38]. To reconstruct SAB MAGs, bins with completeness >50%, contamination <15%, and either a single 16S rRNA sequence matching a SAB OTU or no detectable 16S sequences were selected. SAB bins generated by different binners and classified identically by GTDB-Tk [39] were collected for refinement using CORITES (see **Extended Methods** for the algorithm details), which is available on GitHub (https://github.com/sutormin94/CORITES).

The resulting SAB MAGs and medium-to-high-quality metagenomic bins of non-sponge-associated bacteria, obtained using binny from seawater and SAM metagenomes (2016 and 2018 samples), were used for annotation and further analysis.

### Functional annotation and phylogenetic reconstruction of MAGs

ORFs in the obtained metagenomic bins and reconstructed MAGs were annotated with MetaGeneMark2 v. 1.23 [40] and Prokka v. 1.14.6 [41]. Protein-coding sequences were annotated against the KEGG database using BlastKOALA v. 3.0 [42], anvi’o v. 8 [43] and eggNOG-mapper online resource [44]. Amino acid biosynthetic pathways were predicted with BlastKOALA and GapMind [45]. Biosynthetic gene clusters (BGCs) were predicted with antiSMASH v. 7.0 [46]. Secretion systems were detected with TXSScan (Galaxy Version 2.0 + galaxy3) [47], ELPs were identified using InterProScan v. 5.64-96 [48] with the Pfam database [49]. Signal peptides were predicted with SignalP6.0 [50]. CRISPR-Cas systems were annotated using CRISPRCasTyper [51]. Genetic cluster similarity was analyzed and visualized with clinker [52]. For predicting and comparing proteins structures we used AlphaFold 3 [53] Dali [54].

Full-length 16S rRNA sequences derived from SAB MAGs were searched using blastn against NCBI nt, NCBI 16S rRNA, and SILVA NR99. Sequences retrieved from these databases were combined and clustered with MMseqs2 to remove duplicated sequences. Representative sequences were aligned using MUSCLE in MEGA-X [55] and a maximum likelihood tree was constructed based on the multiple alignments with 100 bootstrap iterations. A bootstrap consensus tree was visualized in iTOL [56].

Refined SAB MAGs were classified using the GTDB-Tk. A tree constructed based on the concatenated alignment of 120 bacterial phylogenetically informative marker proteins (Bac120) was obtained and visualized in iTOL.

### Annotation of sponge transcriptomes

ORFs were predicted in the published assembled transcriptomes for HP (labeled, respectively, Tr1 and Tr2) [57, 58] and *Isodictya sp.* [27] with GeneMarkS-T v. 5.1 [59] and unique sequences longer than 100 amino acids were selected for further analysis. Metabolic pathways were reconstructed using BlastKOALA and KEGG Mapper [60]. To detect the expression of bacterial genes of sponge-associated bacteria, genes from OTU4 and OTU23 MAGs were blasted against HP transcriptomes. Hits were selected if identity and coverage exceeded the 95% threshold.

### FISH microscopy

FISH was performed according to a standard protocol [61] with Cy3-labeled SAB-specific and Cy5-labeled bacterial universal probes [62] for 16S rRNA (**Supplementary Table S2**). Briefly (see a detailed protocol in **Supplementary Methods**), sponge tissue samples were fixed in formaldehyde, dehydrated, and stored at -20°C until further processing. Rehydrated fragments were hybridized with an equimolar mixture of universal and particular SAB-specific probes. DNA was stained with Hoechst-33342. Stained samples were mounted with Prolong Gold antifade (Invitrogen) and imaged in the Airyscan mode [63] using a Zeiss LSM 800 laser scanning confocal microscope at the Center of N.K. Koltsov RAS. Images were processed in ZenBlack (Carl Zeiss), followed by analysis in ImageJ Fiji [64]. Specificity of SAB-specific probes was confirmed by hybridization with tissues from non-cognate sponge species, which showed no detectable signal.

## Results

### Sympatric sponges harbor species-specific sets of host-associated OTUs

In contrast to tropical sponge species, the composition and temporal dynamics of microbiomes associated with Arctic sponges remain understudied. We selected three widely abundant sponge species from the White Sea and sampled them over a period of six years to track changes in the associated microbial taxa and see if ecologically similar sponges harbor similar or distinct microbiomes.

Selected species represent sympatric cold-water marine sponges belonging to Demospongiae/Heteroscleromorpha - *Halichondria panicea* (HP), *Halichondria sitiens* (HS), and *Isodictya palmata* (IP) (**Figure 1A**). Samples were collected in the Kandalaksha Bay of the White Sea in August-September of 2016, 2018, and 2022 along with seawater samples from the collection sites. Metagenomic DNA was extracted from enriched microbiome fractions and subjected to 16S rRNA V3-V4 amplicon sequencing (**Supplementary Table S3**). To investigate the geographical variability of HP microbiomes, we also collected sponges from Dalnye Zelentsy (the Barents Sea) (**Figure 1B**). Additionally, we re-analyzed publicly available metagenomic datasets for HP, *Isodictya kerguelenensis*, and *Isodictya erinacea* from various geographical locations in the Northern and Southern hemispheres (**Supplementary Table S3**).

**Figure 1.**
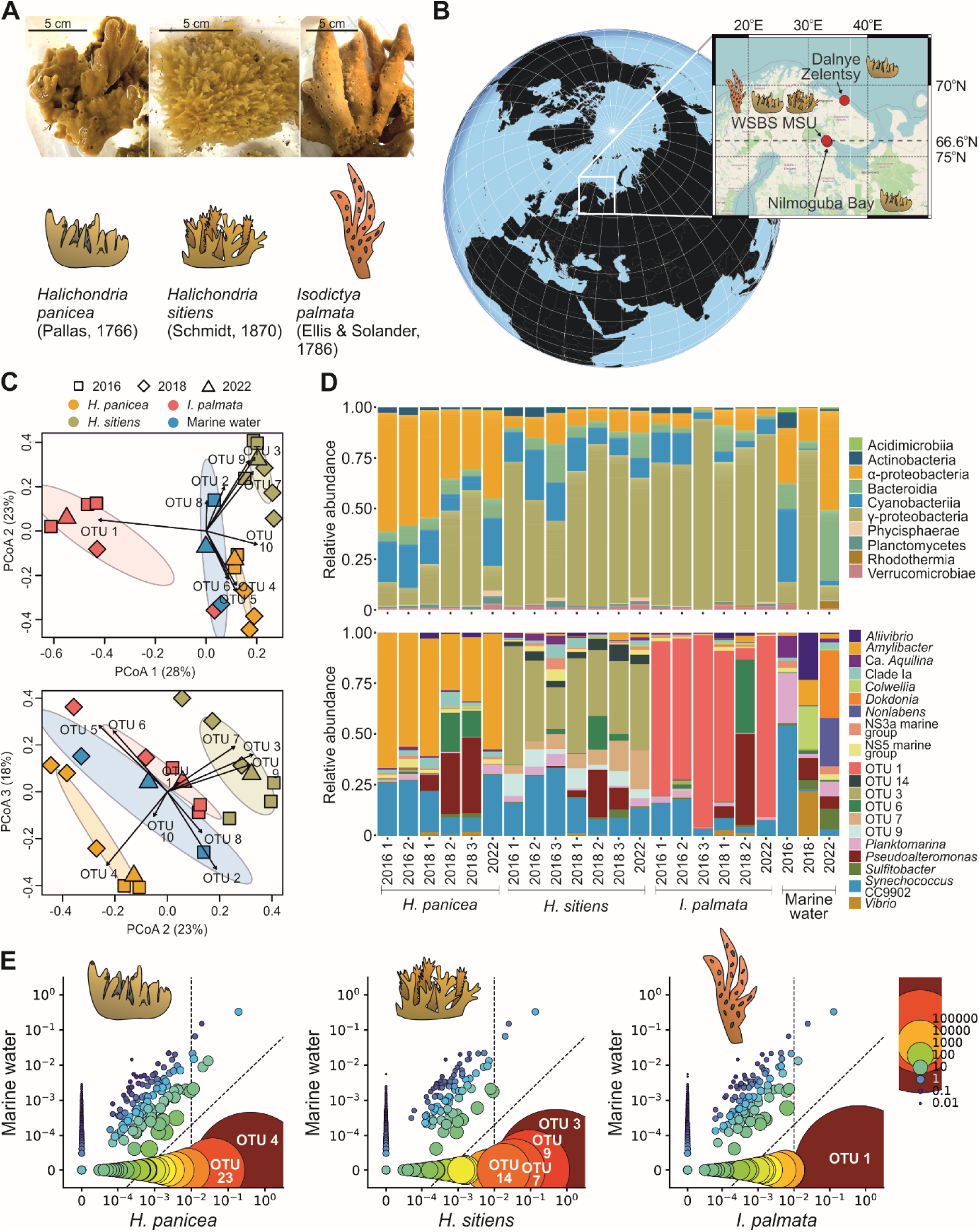
Microbiomes of HP, HS, and IP. (**A**) Representative images of three Porifera species studied. Scale bars of 5 cm are shown. Cartoon icons illustrating the sponge species are shown below the corresponding images. The icons are used throughout the manuscript as organism labels. (**B**) A map of the sample collection sites. (**C**) PCoA of Bray-Curtis dissimilarity between sponge and seawater microbiomes from the White Sea. (**D**) Relative abundance of bacterial classes (top) and genera (bottom) in sponge and seawater microbiomes from the White Sea. (**E**) Symbiont-plots demonstrating identification of SAB OTUs in the sponge microbiomes (data for representative replicates 2016 2 for HP; 2016 1 for HS, and 2016 1 for IP are shown). Each OTU represents a point. The radius and color of each point are proportional to the ratio of relative abundances of the OTU in sponge and seawater. The axes are presented on a logarithmic scale. The vertical dashed line marks a relative abundance of 1% in the sponge microbiome; the diagonal dashed line represents a 50:1 ratio between relative abundances in the sponge and water microbiomes.

Beta-diversity analysis revealed that the studied sponges harbored distinct SAMs, which significantly differed in composition from the surrounding seawater (**Figure 1C**, **Supplementary Figure 1A**). SAMs had comparable numbers of operational taxonomic units (OTUs) with seawater but lower Shannon and higher Simpson indices (**Supplementary Figure 2**) suggesting the presence of highly represented sponge-specific OTUs (**Figure 1D**). This pattern indicates that the studied sponges belong to the LMA group [65, 66].

Indeed, the microbiome of HP from the White Sea was dominated by genus *Amylibacter* OTU4 (**Figure 1D**), sharing 100% identity with 16S from a HP symbiont *Ca.* Halichondribacter symbioticus (**Supplementary Table S4**) [67]. Similarly, it dominated microbiomes of HP collected from geographically diverse locations, implying a tight symbiotic association with its host (**Table 1**) [68, 69]. The HS microbiome was dominated by OTU3 which belongs to the UBA10353 order (Gammaproteobacteria) (**Figure 1D**, **Table 1**, **Supplementary Table S4**). The OTU1 was dramatically overrepresented in the IP, except for one sponge collected in 2018 (**Figure 1D**, **Table 1**). The OTU1 sequence perfectly matched one obtained from the sponge *Haliclona sp*. and had 98% and 96% identity with dominant OTUs from the microbiomes of Antarctic sponges *I. kerguelenensis* [70] and *I. erinacea* [18], which may suggest symbiotic association with sponges.

**Table 1.**
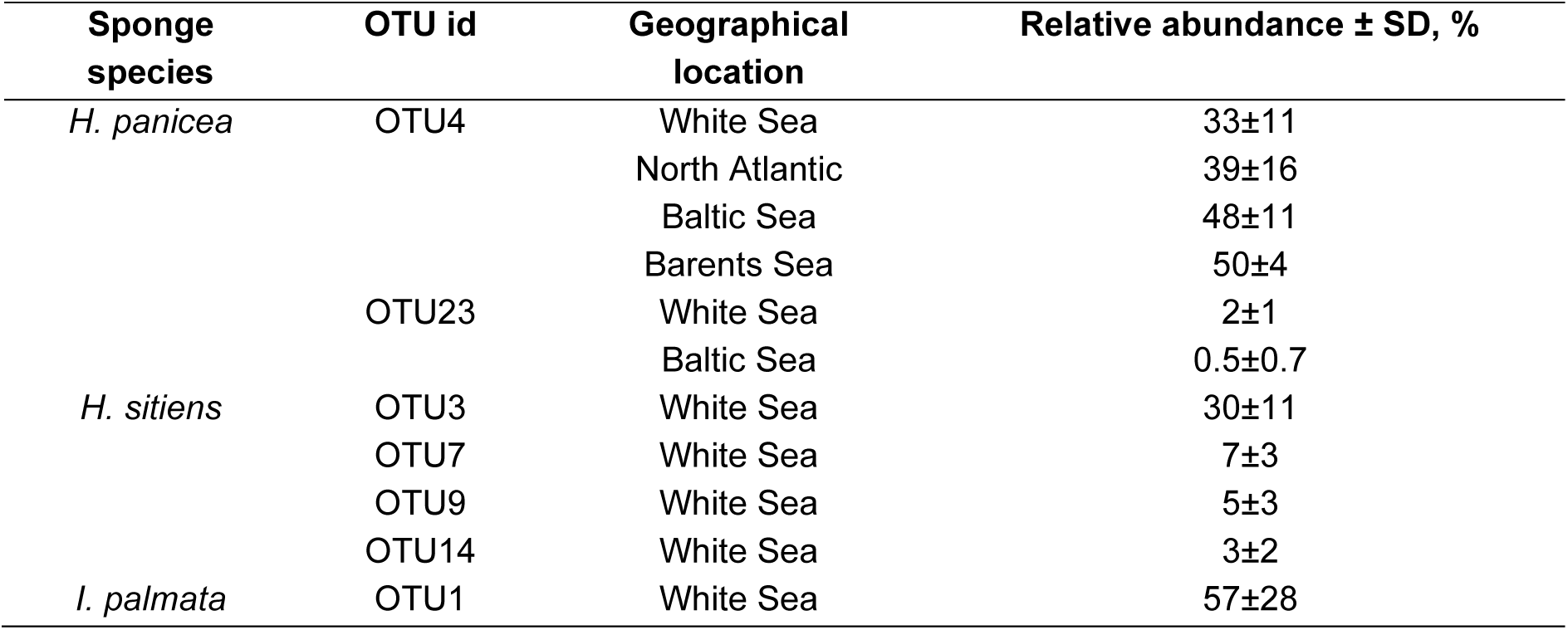
Relative abundance of identified OTUs associated with sponges.

After determining the dominant species of the microbiome, we aimed to identify low-abundant SABs. Using the criteria for low-abundant SABs (see **Methods**), we found OTU23 (Puniceispirillales, Alphaproteobacteria), which was frequently associated with HP microbiomes and was absent in seawater (**Figure 1E**, **Table 1**, **Supplementary Tables S4-5**). OTU23 was found in nearly half of the HP individuals collected from the Baltic Sea (14/26 samples) [69] and was also identified with Pebblescout [71] in shotgun metagenomes of HP collected in Scotland. Interestingly, it was scarce in samples from the North Atlantic (14/84 samples) and the Barents Sea (0/2 samples), indicating that OTU23 likely is not an essential member of the HP microbiome but is frequently associated with this sponge. Similarly, we detected three additional OTUs associated with HS sponges (**Figure 1E**, **Table 1**) - OTU7 (Gammaproteobacteria, Pseudomonadales), OTU9 (Gammaproteobacteria, HOC36), and OTU14 (Gammaproteobacteria), two of which, OTU9 and OTU14 shared a high level of identity with 16S sequences obtained from other sponge species (**Supplementary Tables S4**). No additional OTUs were found to be associated with IP (**Figure 1E**).

A consecutive sampling of sponge microbiomes from the WSBS location allowed us to observe their temporal dynamics. Identified SAB OTUs were stably associated with sponge species over the 6-year period (**Figure 1D**, **Supplementary Figure 1A**). However, in the 2018 data, we observed that several SAB OTUs (OTU1, OTU3, OTU14, and OTU23) were also detected at low levels in seawater (<0.01% relative abundance), whereas OTU4 reached a relative abundance of 8%. In non-host sponges, these OTUs were observed at low levels (mean <0.15% relative abundance) across all sampling timepoints (**Supplementary Figure 1A**). Despite bacterial exchange between sympatric sponges, no new prolific interactions were established, indicating a specific symbiosis.

Therefore, our 16S metagenomic data suggest that the studied sympatric Arctic sponge species harbor distinct SAMs, dominated by strongly host-associated species-specific OTUs, which likely represent bacterial symbionts.

### Recovery of SAB MAGs and their refining with CORITES

To reconstruct SAB genomes, we sequenced sponge metagenomes with short- and long-read technologies and obtained initial metagenomic bins using several commonly used binners. Bins obtained from long-read assemblies were more continuous, however, they typically contained several different 16S and/or 23S sequences, indicating contamination. To avoid possible chimeric sequences, long-read assemblies were excluded from further analysis.

Multiple NGS datasets generated over the 6-year sampling period allowed us to obtain numerous bins for identified SABs. Phylogeny reconstruction using GTDB-Tk demonstrated clustering of related bins, indicating a robust classification independently of the dataset and/or binning algorithm used (**Supplementary Figure 3**). Due to the differences between binning approaches, related bins obtained with different binners from the same metagenome had a considerable fraction of unshared contigs (on average, 62±28% contigs were shared). To reduce the frequency of binning errors, refine and scaffold bins, we developed and applied the CORITES algorithm (**Supplementary Note 1**). Using CORITES, we were able to increase completeness and reduce contamination for most sponge-associated MAGs compared to the initial bins (**Supplementary Table S6**). Reconstructed MAGs have high completeness (>80%, except for the OTU23 from HP), low contamination (<5%), and nearly complete sets of tRNA genes indicating their medium-to-high quality by MIMAG criteria [72] (**Supplementary Table S7**). A low completeness level (63%) and small size of the OTU23 MAG can indicate incomplete assembly or genome reduction due to a potential symbiotic lifestyle.

Overall, we were able to reconstruct MAGs for all SABs identified with 16S metagenomics.

### Localization of SABs in sponge tissues

To orthogonally confirm the presence of identified SABs in sponge tissues and study their localization, we performed FISH microscopy using bacteria-specific oligonucleotides for sponge tissue sampled from different body sites – the osculum region, the middle and the bottom sections of the sponge body. Cy3-labeled probes (red) was used for selected bacteria and Cy5-labeled probes (green) for all bacteria.

We found that SABs were present in all sampled body sites. Cells of the major HP SAB, OTU4, were small and rod-shaped, matching the morphology described previously for the *Ca*. H. symbioticus [67]. The bacteria were presumably accumulated within some host cells (bacteriocytes) and also found scattered in mesohyl (**Figure 2A**). We were unable to obtain a reliable image of SAB OTU23 (HP), likely due to its low abundance in the sponge (2±1% based on 16S data). HS tissues were densely populated with SABs. The dominant SAB of HS, OTU3, had rod-shaped cells (**Figure 2B**). Other SABs, OTU7 and OTU9, also displayed a rod-shaped morphology, while OTU14 cells were crescent-shaped (**Figure 2D-F**). Cells of OTUs 3, 9, and 14 were supposedly associated with bacteriocytes and located as individual cells intercellularly, while OTU7 cells were primarily intercellular. A single SAB from IP, OTU1, dominated in tissue samples in accordance with 16S data. The OTU1 cells exhibited spherical morphology and were likely clustered in intercellular microcolonies (**Figure 2C**).

**Figure 2.**
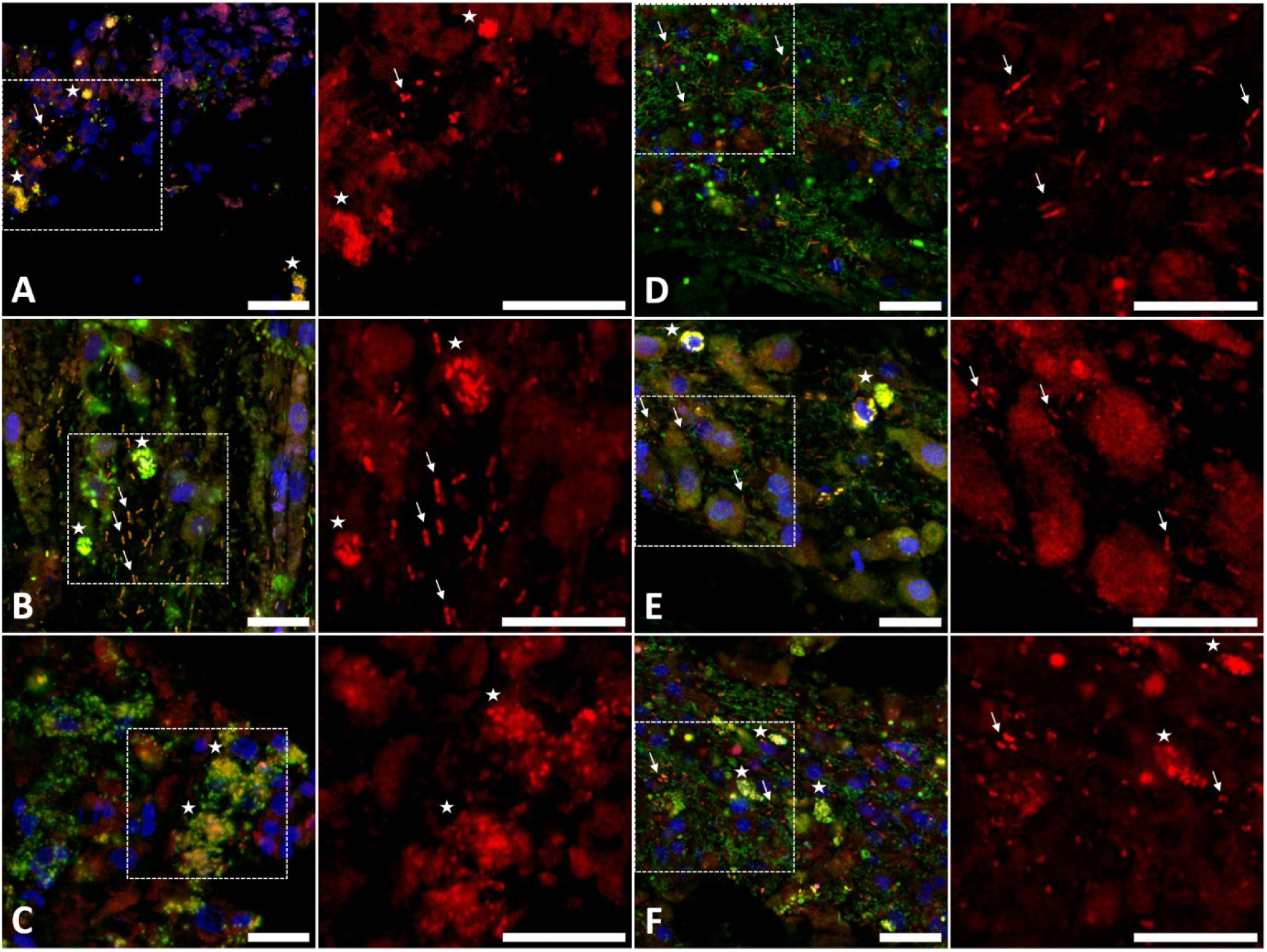
FISH images of bacteria in tissue sections of the studied sponges. Bacteria were labeled with SABs-specific Cy3 (red) and universal probes with Cy5 (green). DNA was stained with Hoechst-33342 (blue). (**A**) Section of HP hybridized with a probe specific for OTU4, (**B**) Section of HS hybridized with a probe specific for OTU3, (**C**) Section of IP hybridized with a probe specific for OTU1, (**D**) Section of HS hybridized with a probe specific for OTU7, (**E**) Section of HS hybridized with a probe specific for OTU9, (**F**) Section of HS hybridized with a probe specific for OTU14. For all panels, a merge of three color channels is shown on the left; enlarged field indicated by a white rectangle in the left image is shown on the right in red (Cy3) channel only. Arrows and asterisks indicate individual cells and their clusters, respectively. Scale bars: 10 μm.

Taken together, we localized all identified SABs (except for OTU23) in different body sites of sponges and found that they exhibit distinct patterns of organization in tissues.

### General metabolic characteristics of the reconstructed sponge-associated MAGs

To gain insights into the functional potential of SABs and detect possible metabolic interactions within a holobiont, we analyzed their metabolic capabilities. By reconstructing putative pathways in sponge-associated MAGs using KEGG, we identified nearly complete glycolysis and tricarboxylic acid cycle pathways present in most of them. OTU14 MAG (HS), however, was lacking the pyruvate dehydrogenase complex (M00307), tricarboxylic acid cycle, and its variation, the glyoxylate cycle, possibly indicating genomic reduction (**Figure 3**). An incomplete pentose phosphate pathway was detected in several MAGs from different sponges. Autotrophic carbon fixation was not reliably identified in any of MAGs. Although the completeness of the reductive pentose phosphate cycle and the reductive citrate cycle in some genomes, such as *Ca.* H. symbioticus from HP (see **Supplementary Note 2**), reached 70%, the same genes are also involved in oxidative pathways. Therefore, we inferred that all identified SABs are heterotrophic. SAB MAGs also carried genes of cytochrome *c* oxidase, other components of the respiratory electron transport chain, and ATP-synthase, likely utilized for ATP production (**Figure 3**). No nitrogen metabolic pathways were detected in MAGs [73].

**Figure 3.**
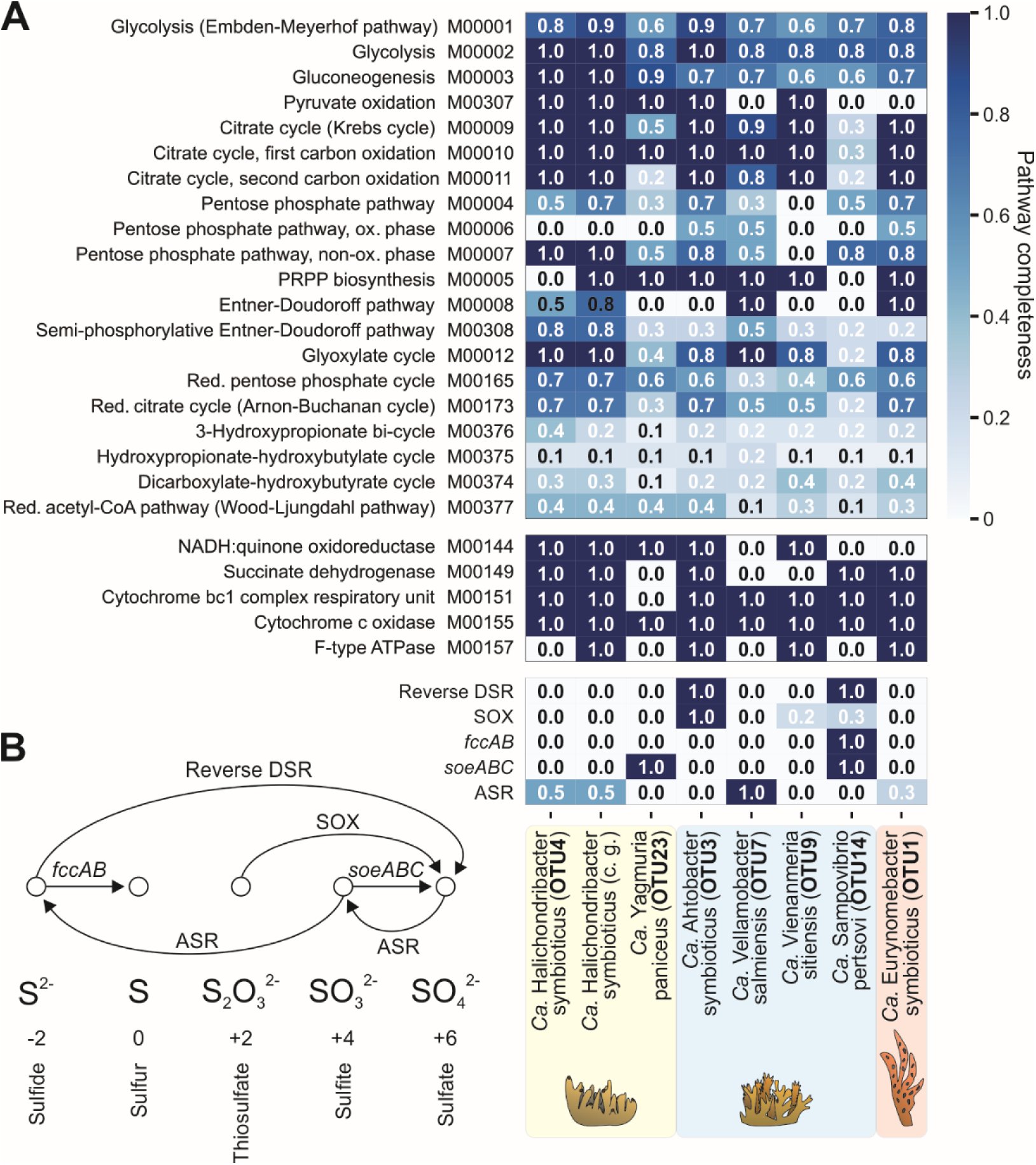
General carbohydrate, energy, and sulfur metabolic pathways identified in SAB MAGs. (**A**) Heatmap representation of the completeness of metabolic pathways. Heatmap values indicate the completeness of gene sets ranging from 0 (no genes of a pathway detected) to 1 (a complete pathway detected), based on KEGG annotation. The OY365741.1 genome, identified as *Ca*. H. symbioticus, was included in the analysis. (**B**) A scheme of common sulfur transformation pathways. ASR – assimilatory sulfate reduction; DSR – dissimilatory sulfate reduction; SOX –sulfur oxidation system; c.g. – complete genome.

We detected diverse sulfur metabolic pathways enriched in SAB MAGs from HS. OTU3 MAG encoded the *dsr* operon (*dsrABLEFHCMKJOP* genes) which is likely responsible for oxidation of sulfide to sulfite via the reverse dissimilatory sulfate reduction (DSR) pathway as it lacked the *dsrD* gene and included *dsrEFH* genes [74]. Additionally, this MAG contained genes of the sulfur oxidation (SOX) system [75] - the *soxABXYZ* operon responsible for oxidation of thiosulfate to sulfate - and lacked *soxCD* genes (**Figure 3**). The presence of both DSR and SOX pathways suggests that OTU3 bacterium may utilize sulfide and thiosulfate as electron donors, as was hypothesized for two unrelated sponge symbionts with a similar composition of pathways [76, 77]. The genome of OTU14 MAG encoded a complete reverse-DSR sulfur oxidation pathway and carried *soeABC* genes for oxidation of sulfite to sulfate [78], flavocytochrome-C dehydrogenase (*fccAB* genes) for sulfide oxidation to elemental sulfur [79], and a partial *sox* operon (*soxXY* genes) (**Figure 3**). OTU9 MAG also contained a partial *sox* operon (*soxCDY*) which may be attributed to incomplete genome assembly. In contrast to other HS SABs, OTU7 MAG lacked pathways of sulfur oxidation but had a complete assimilatory sulfate reduction (ASR) pathway, reducing sulfate to sulfite and further to sulfide [80]. Among bacteria from other sponges, only OTU23 MAG (HP) had genes related to sulfur metabolism and contained the *soeABC* operon (**Figure 3**).

The enrichment of sulfur transformation pathways, particularly oxidation, in HS SABs suggests specialization of this community in sulfur metabolism and its potential role in sulfur cycling within the holobiont.

### SABs may complement host metabolism

To identify metabolic capabilities, absent in host sponges and potentially complemented by SABs, we analyzed publicly available transcriptomes for HP (Tr2) and *Isodictya* sp (**Supplementary Note 3**). No transcriptomic data were published for HS, but considering that it is closely related to HP (99.6% identity by 18S), we assumed that they could have similar metabolic potential.

We analyzed the biosynthetic pathways of proteinogenic amino acids in the HP and *Isodictya sp.* transcriptomes and found no expression of pathways for branched-chain (Ile, Leu, Val), non-polar (Met), polar (Thr), positively charged (Arg, Lys), and aromatic (His, Trp, Phe, Tyr) amino acids pathways, suggesting that these amino acids are essential for sponges. At the same time, these nearly complete pathways were identified in MAG of at least one SAB from each of the three studied sponge species, except for the Phe/Tyr biosynthetic pathways (**Figure 4A**). Further prediction of Phe/Tyr biosynthetic pathways in SAB MAGs using GapMind revealed nearly complete pathways in HS SABs (OTU3, OTU7, OTU14) and in the IP SAB OTU1. Moreover, this pathway was also found in the *Ca*. H. symbioticus genomes (OY365741.1 and OY365738.1) suggesting that its absence in the OTU4 MAG from HP was due to genome incompleteness.

**Figure 4.**
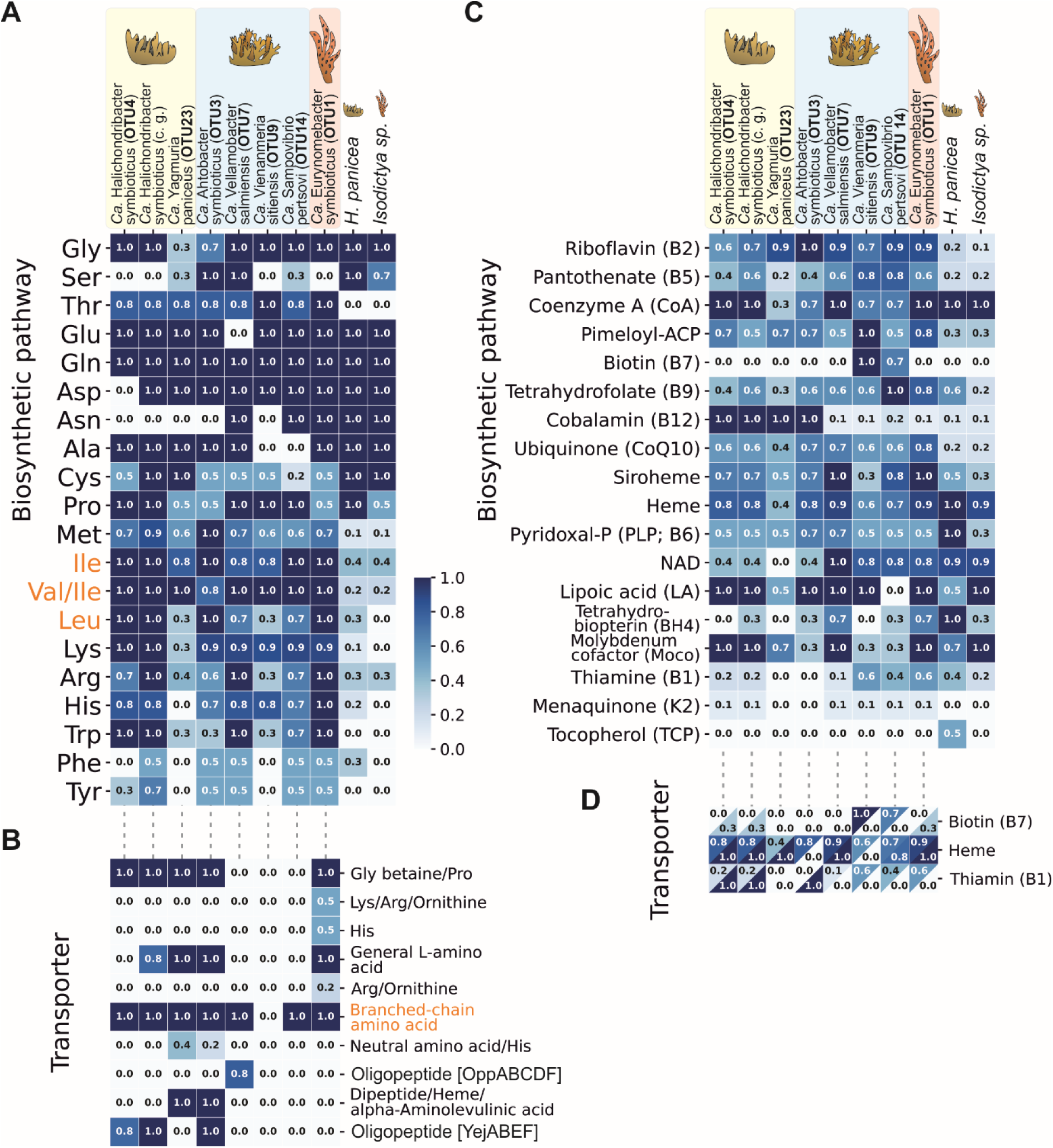
*De novo* amino acid (**A**) and vitamin/cofactor (**C**) biosynthetic pathways identified in SABs and sponge transcriptomes. Transporters related to amino acid (**B**) and vitamin/cofactor (**D**) import/export. Heatmap values represent the completeness of gene sets from 0 (no genes of a pathway/transporter detected) to 1 (a complete pathway/transporter observed), based on KEGG annotation. In panel D, the heatmap values in the upper and lower triangles represent, respectively, the completeness of biosynthetic pathways for vitamins and their corresponding transporters.

Next, we analyzed the presence of genes encoding amino acid and peptide transporters in SAB MAGs. SAB MAGs from all three microbiomes harbored genes for a general L-amino acid ABC transporter, and all SAB MAGs, except OTU9 (HS), encoded an ABC transporter for branched-chain amino acids (**Figure 4B**). Moreover, the expression of branched-chain amino acid and general L-amino acid transporters was detected for both OTU4 and OTU23 SABs in the HP Tr1 transcriptome. Since ABC transporters can function bidirectionally [81, 82], SABs may potentially supplement their hosts with essential amino acids (particularly branched-chain ones). Taken together, our analysis suggests a potential complementation of sponge requirements in essential amino acids by SABs.

Interestingly, a bacteria consortium in HS showed a division of biosynthetic functions: complete Met and Leu pathways were found exclusively in OTU3 MAG, while Arg and Trp pathways were present only in OTU7 MAG. OTU9 MAG was the only one lacking Phe and Tyr pathways, while nearly complete pathways for other amino acids (Ile, Val, Lys, His, Thr) were present in all four MAGs. This specialization may explain the relatively complex community in HS and suggests potential cross-feeding among its SABs. In contrast, OTU1 MAG, the sole SAB form IP, has nearly complete or complete pathways for all the above-mentioned presumably essential amino acids indicating that it can likely fulfill the metabolic need of its host alone.

Similarly, we analyzed the potential complementation of biosynthetic pathways of vitamins and cofactors between MAGs and sponge transcriptomes. Biosynthetic pathways of several B vitamins (B1, B2, B5, B7, B12), pimeloyl-ACP, ubiquinone, and siroheme were scarce in both sponge transcriptomes. Among them, nearly complete or more complete, if compared to sponge, pathways for B2, B5, pimeloyl-ACP, ubiquinone, and siroheme were found in a MAG of at least one SAB from each of the three studied sponge species (**Figure 4C**). Pathways for B1 were identified in OTU1 MAG (IP) and OTU9 (HS), but not in MAGs from HP, suggesting an alternative source of this vitamin for this sponge species. Similarly, B12 pathways were only present in SAB MAGs from HP and HS, but not in OTU1 MAG from IP, while B7 pathways were exclusively found in SAB MAGs from HS and not in IP or HP SAB MAGs. Menaquinone and tocopherol pathways were absent in all SAB MAGs and sponge transcriptomes and thus are likely acquired from a different source by the holobionts. Analysis of vitamin/cofactor transporters revealed a counter-association with their biosynthetic pathways for B7 and, especially, for B1 (**Figure 4D**), which suggests that auxotrophic SABs encode transporters to obtain compounds from the environment (presumably, from other members of the microbiome).

Our analysis indicates that SABs can potentially complement the metabolism of host sponges with all essential amino acids and multiple vitamins.

### Taurine dissimilation pathways are present in genomes of major SABs

To establish stable symbiotic relationships, the flow of metabolites might be reciprocal, i.e., symbiotic bacteria should gain from feeding on sponge-produced compounds. In a search of such metabolic exchanges, we focused on taurine, a prevalent compound found in sponge tissues, though sponges have not yet been experimentally demonstrated to synthesize and release it [83, 84]. Dissolved taurine is an important source of carbon and energy for seawater prokaryotic communities [85, 86], and similarly might be crucial for sponge symbionts [14].

Previously, taurine biosynthetic pathways were identified in the complete genome of the sponge *Amphimedon queenslandica* [87]. Using KEGG, we revealed complete pathways for taurine biosynthesis from cysteine in HP (Tr2) and *Isodictya sp*. transcriptomes, supporting the potential host-derived origin of this compound (**Figure 5A**). In contrast, all SAB MAGs lacked taurine biosynthetic pathways.

**Figure 5.**
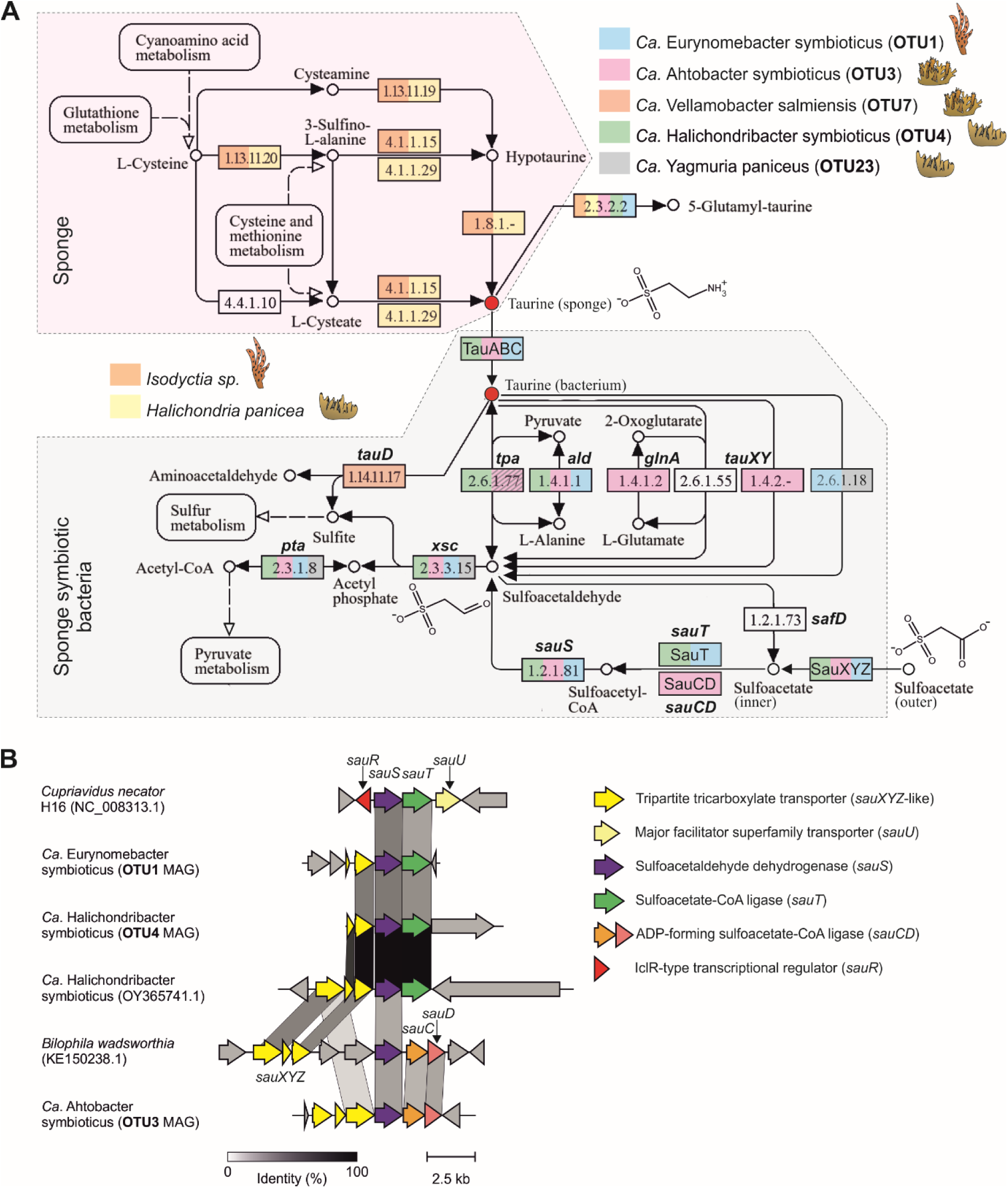
Taurine and sulfoacetate metabolism pathways in sponges and SABs (**A**) Metabolic map of taurine and sulfoacetate biosynthesis and degradation according to KEGG. The shading of the *tpa* gene in the OTU3 MAG indicates that this gene was found in several original OTU3 metagenomic bins but was absent in the final OTU3 MAG. (**B**) Comparison of characterized gene clusters involved in degradation of sulfoacetate from *B. wadsworthia* and *C. necator* and putative gene clusters from major SABs.

We found that MAGs of all dominant SABs (OTU4, OTU3, and OTU1) from all three sponge species encoded putative taurine ABC transporters (*tauABC*), thus suggesting active import of taurine. Among these bacteria, OTU4 (HP) and OTU3 (HS) MAGs harbored genes encoding enzymes involved in taurine catabolism, converting it into sulfite and acetyl-CoA via sulfoacetaldehyde and acetyl-phosphate intermediates: taurine:pyruvate aminotransferase (*tpa,* the gene was found expressed in the HP Tr1 transcriptome), sulfoacetaldehyde acetyltransferase (*xsc*), and phosphate acetyltransferase (*pta*). OTU1 MAG (IP) and phylogenetically related bacteria carried *xsc* and *pta*, but lacked the *tpa* gene (**Supplementary Table S9**, **Figure 5A**). However, they encoded a ω-amino acid aminotransferase (EC 2.6.1.18) that converts taurine into sulfoacetaldehyde, thus substituting for Tpa activity [88, 89] (**Supplementary Table S9**, **Figure 5A**). Additionally, we revealed metabolic genes from other pathways associated with taurine catabolism in MAGs of dominant SABs (**Supplementary Note 4**). Notably, a complete pathway, including the *tauABC*, *tpa*, *xsc*, and *pta* genes, was also identified in 8 of 99 non-sponge-associated metagenomic bins retrieved from seawater and SAMs, all of which belonged to Alphaproteobacteria (**Supplementary Table S9**).

Genomes of minor SABs lacked genes encoding taurine transporters. OTU7 MAG (HS) carried the gene for taurine dioxygenase (*tauD*) responsible for taurine conversion to sulfite and aminoacetaldehyde [90], while OTU23 MAG (HP) carried a complete pathway with ω-amino acid aminotransferase, *xsc*, and *pta* genes present (**Figure 5B**).

Our data indicate that taurine catabolism is widespread among taxonomically unrelated SABs, which suggest that this compound can be potentially utilized by SABs as a source of carbon, sulfur, and nitrogen.

Given that taurine may serve as an important nutrient source for SABs, we attempted to cultivate major SABs from HP and HS using various media, including poor and rich compositions supplemented with taurine (see **Supplementary Methods**). However, we did not obtain any colonies corresponding to OTU4 or OTU3 bacteria at any conditions, suggesting that taurine alone is not the limiting factor for their growth and that these bacterial residents are strongly associated with a host-specific environment.

### Major SABs encode putative gene clusters responsible for uptake and degradation of taurine derivative, sulfoacetate

Another compound linked to taurine metabolism is sulfoacetate, which can be produced by deamination and oxidation of taurine by some marine bacteria during the assimilation of taurine nitrogen [91]. This compound can also be imported and metabolized by other bacteria via the sulfoacetaldehyde pathway, allowing it to be used as a source of carbon and energy [92–94].

We noticed that all three dominant SABs encoded genes involved in the conversion of sulfoacetate to sulfoacetaldehyde (**Figure 5A**). OTU4 (HP) and OTU1 (IP) MAGs carried gene clusters resembling the *sauST* cluster from *Cupriavidus necator* H16 and encoded sulfoacetaldehyde dehydrogenase (SauS) and sulfoacetate-CoA ligase (SauT) [93]. OTU3 MAG (HS) contained an alternative gene cluster with the *sauT* gene substituted with *sauCD* (encoding the ADP-forming sulfoacetate-CoA ligase) genes, resembling a *sauSCD* cluster responsible for sulfoacetate processing in *Bilophila wadsworthia* [94]. All detected *sau* gene clusters were associated with genes of tripartite tricarboxylate transporter (TTT) encoding a putative sulfoacetate transporter SauXYZ previously identified in the *B. wadsworthia*. The putative *sauST* gene clusters from OTU1 and OTU4 MAGs therefore comprised a novel hybrid type associated with a TTT transporter instead of a single-gene-encoded SauU MFS transporter described in *C. necator* [93] (**Figure 5B**).

Sulfoacetate metabolism genes were rare in non-sponge-associated metagenomic bins: *sauST* and *sauSCD* clusters were detected in 4/99 and 3/99 of such bins, respectively. Interestingly, 4 of these bins belong to the same orders as SABs from HP (Puniceispirillales or Rhodobacterales) (**Supplementary Table S9**).

Although we did not detect the previously described *safD* pathway responsible for sulfoacetate production in a marine bacterium [91] within the studied SABs or other metagenomic bins from SAMs, we hypothesize alternative pathways exist within a holobiont and suggest that SABs may utilize the released sulfoacetate as an additional carbon and energy source.

### Genomes of SABs encode symbiosis-associated genes

Next, we focused on genes known to be associated with symbiotic lifestyle. For example, bacterial eukaryotic-like proteins (ELPs) are considered to be involved in symbiont-host interactions [15, 95] and were found enriched in sponge metagenomes and SAB genomes [16, 17, 19]. To investigate this association, we predicted ELP-domain-containing proteins in the reconstructed SAB MAGs and other medium-to-high-quality metagenomic bins from sponge and seawater metagenomes (**Supplementary Figure 4**).

Principal component analysis (PCA), performed on ELP domain frequencies, separated SAB MAGs from each other and the majority of non-sponge-associated metagenomic bins, indicating that SAB MAGs had different sets of enriched ELP domains (**Figure 6A**). Indeed, tetratricopeptide repeat domains (TPR) were abundant in OTU1 (IP) and OTU9 (HS) MAGs, as their normalized frequencies were significantly higher compared to 99 control bins (t-test p-value 4.2e-2). Ankyrin domains (Ank) were substantially enriched in OTU1 MAG (t-test p-value 1.7e-3). Fibronectin domains (Fn3) were associated with and significantly enriched in OTU14 MAG (HS) (t-test p-value 3.1e-4). Sel1-like repeats (Sel1) were highly enriched in the rest (5 out of 7) of SAB MAGs (t-test p-value 8.0e-13) (**Figure 6B**, **Supplementary Figure 5A**). Similar associations were observed for frequencies of ELP-encoding genes (**Supplementary Figures 5B**, **C**). A substantial fraction of ELP-containing proteins (46/123 or 37%) from SAB MAGs were predicted to have signal peptides for translocation and secretion.

**Figure 6.**
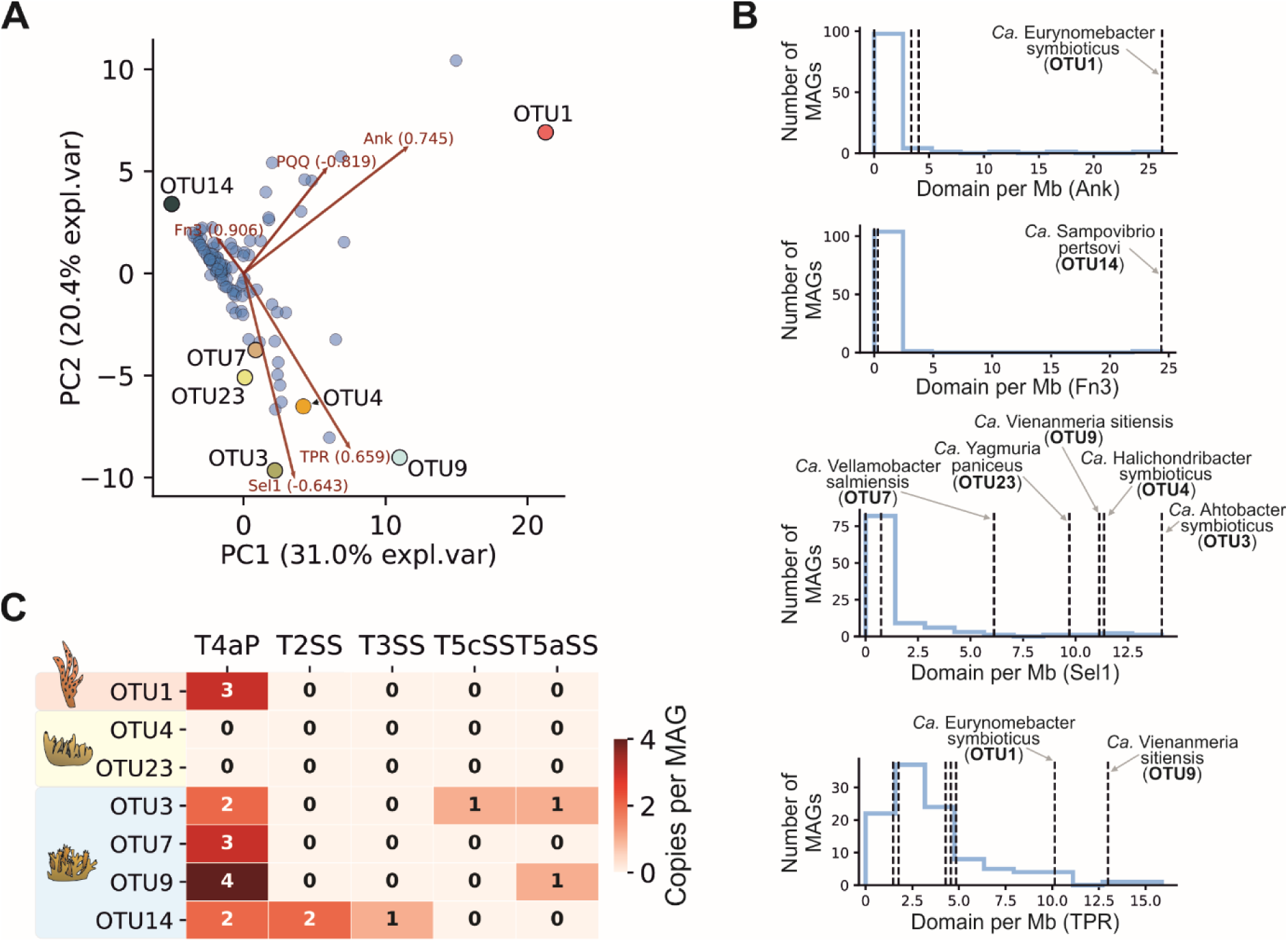
Eukaryotic-like proteins and transport systems identified in SAB MAGs. (**A**) PCA of ELP domain frequencies in SAB MAGs and other metagenomic bins reconstructed from SAM and seawater metagenomes. (**B**) Distributions of normalized ELP domain frequencies in SAB MAGs and other metagenomic bins. Data are shown for ELPs enriched in SAB MAGs. Frequencies of ELPs in SAB MAGs are indicated with vertical dashed lines. (**C**) A heatmap representing the occurrence of secretion system and pili gene clusters in SAB MAGs.

To determine if any of the predicted ELP genes were expressed in *Ca*. H. symbioticus (OTU4 MAG from HP), we re-analyzed the published HP Tr1 (meta)transcriptome (**Supplementary Note 3**) and identified active expression of a Sel1-containing protein with a predicted export signal peptide (Sec/SPI-type, score 0.99). Structural predictions using AlphaFold 3 and a similarity search using Dali revealed a high level of similarity to the exported Sel1-containing protein LpnE from *Legionella pneumophila*, which is required for invasion of eukaryotic cells [96]. These data first confirm the accumulation of certain ELPs in SABs and, second, potentially indicate the role of Sel1 in mediating host-symbiont interactions and the association of *Ca*. H. symbioticus with HP cells.

Another group of pro-symbiotic factors includes secretion systems and other mechanisms involved in interactions with the extracellular matrix. Both bacterial symbionts and pathogens often encode such molecular tools which allow them to interact with host cells (using adhesins, pili, curli, fimbriae) or use secretion systems to export effector proteins that manipulate the host [97, 98]. To study the repertoire of these systems in SAB MAGs, we utilized TXSScan. We observed that type-4 pili (T4aP) were encoded in all SAB MAGs, except HP-associated OTU4 and OTU23 (**Figure 6C**). Interestingly, these two SABs did not encode any of the secretion system-related gene clusters, indicating an alternative mechanism of binding or interaction with host cells (e.g., mediated by exported ELPs). The absence of secretion system genes was unlikely due to incompleteness of the MAGs, at least for *Ca.* H. symbioticus, because its complete genome did not contain any related gene clusters. Bacteria associated with HS encoded unique combinations of gene clusters, likely reflecting the specialization of these bacteria for different niches in the host organism.

Two paralogous genes from the OTU4 MAG (HP), both annotated as ‘Invasion protein B’ (*ialB*), were found to be actively expressed in the HP Tr1 transcriptome. Homologs of these proteins were also identified in OTU23 (HP) and OTU3 (HS) MAGs. Secretory signal sequences (Sec/SPI-type) were predicted in these proteins with a high confidence (>0.97). *IalB* genes were previously detected in the genomes of *Pseudovibrio* bacteria isolated from different marine ecosystems, including the sponge *Polymastia penicillus* and other invertebrates [99, 100]. *IalB* from the human pathogen *Bartonella bacilliformis* is a virulence factor [101] important for the invasion of erythrocytes. The expression of the *ialB* gene by sponge SABs may be important for host-microbe interactions, possibly mediating phagocytosis in sponge cells.

The persistence of identified SABs in sponge microbiomes over a six-year period, combined with the analysis of their MAGs, which revealed several key symbiotic features including the enrichment of ELPs, suggests that the studied SABs are sponge symbionts.

### Identified SABs represent new bacterial taxa of sponge symbiotic bacteria

The phylogenetic analysis of recovered SAB MAGs and their full-length 16S rRNA sequences allowed us to infer their taxonomy.

Both methods unambiguously assigned the HP-dominating OTU4 SAB to *Ca.* H. symbioticus (**Figure 7**). The MAG of the minor HP-associated bacterium, OTU23, was classified as a new member of genus JAGWAQ01 within the UBA1172 family (Puniceispirillales, Alphaproteobacteria) (**Figure 7**). According to 16S sequencing results, OTU23 MAG belonged to a clade of sponge- and coral-associated bacteria known as “Calcibacteria” [102] and to a larger clade which is sister to the Puniceispirillaceae family (**Supplementary Figure 6**). We propose a new name *Ca.* Yagmuria as a replacement for the JAGWAQ01 genus and the species name *Ca.* Yagmuria paniceus for the OTU23 bacterium: the genus name commemorates our colleague Eldar Yagmurov who passed in 2022, and the species name reflects the host name. For the UBA1172 family, we propose a replacement name *Ca.* Calcibacteraceae after the “Calcibacteria” group (**Table 2**).

**Figure 7.**
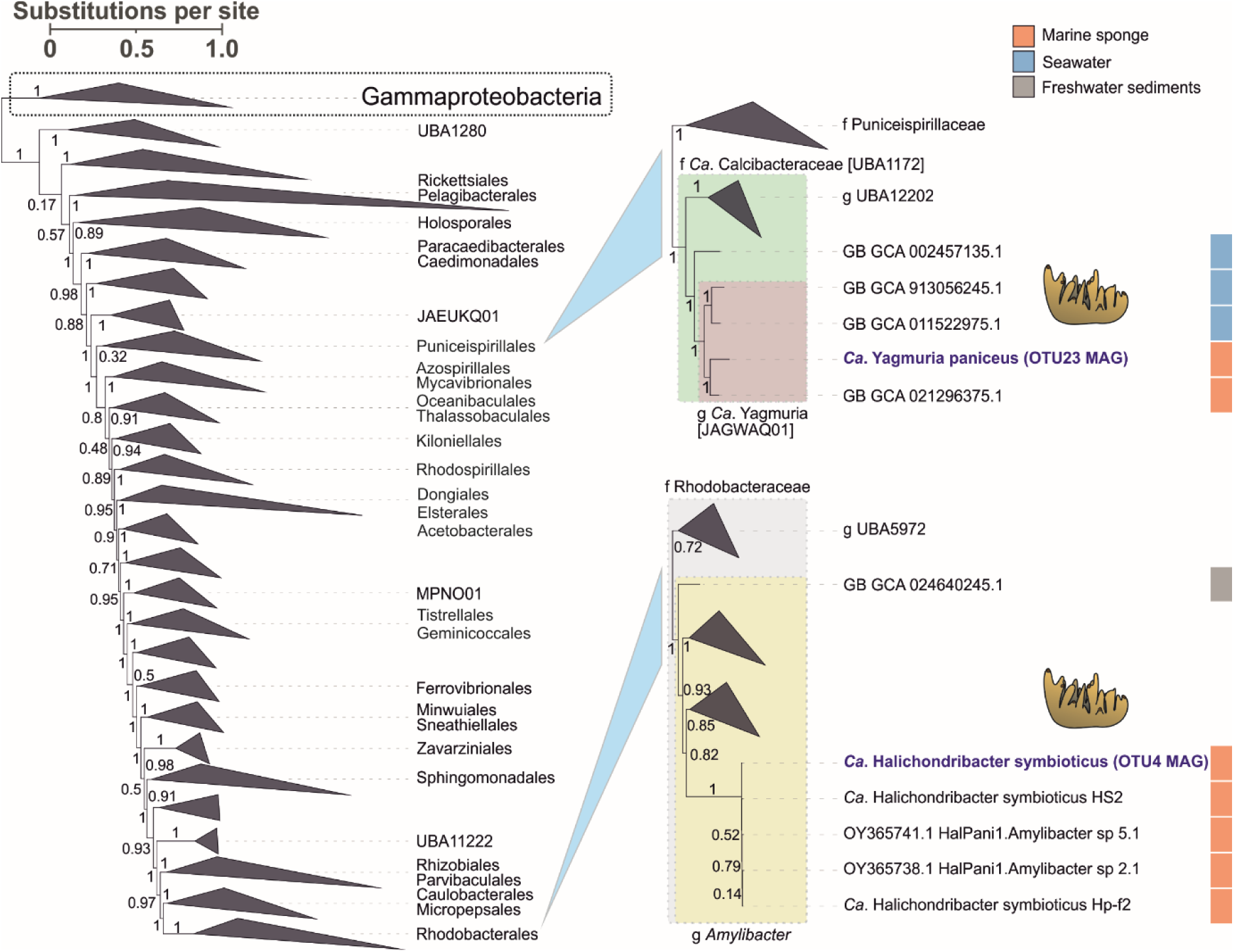
Maximum-likelihood tree of Alphaproteobacteria. The Gammaproteobacteria clade is shown as an outgroup. Local support values obtained with the Shimodaira-Hasegawa test are indicated for nodes (1000 resamples). SAB MAGs obtained in this study are highlighted in blue. Host sponges are indicated with cartoon icons. The isolation source of MAGs is color-coded.

**Table 2.**
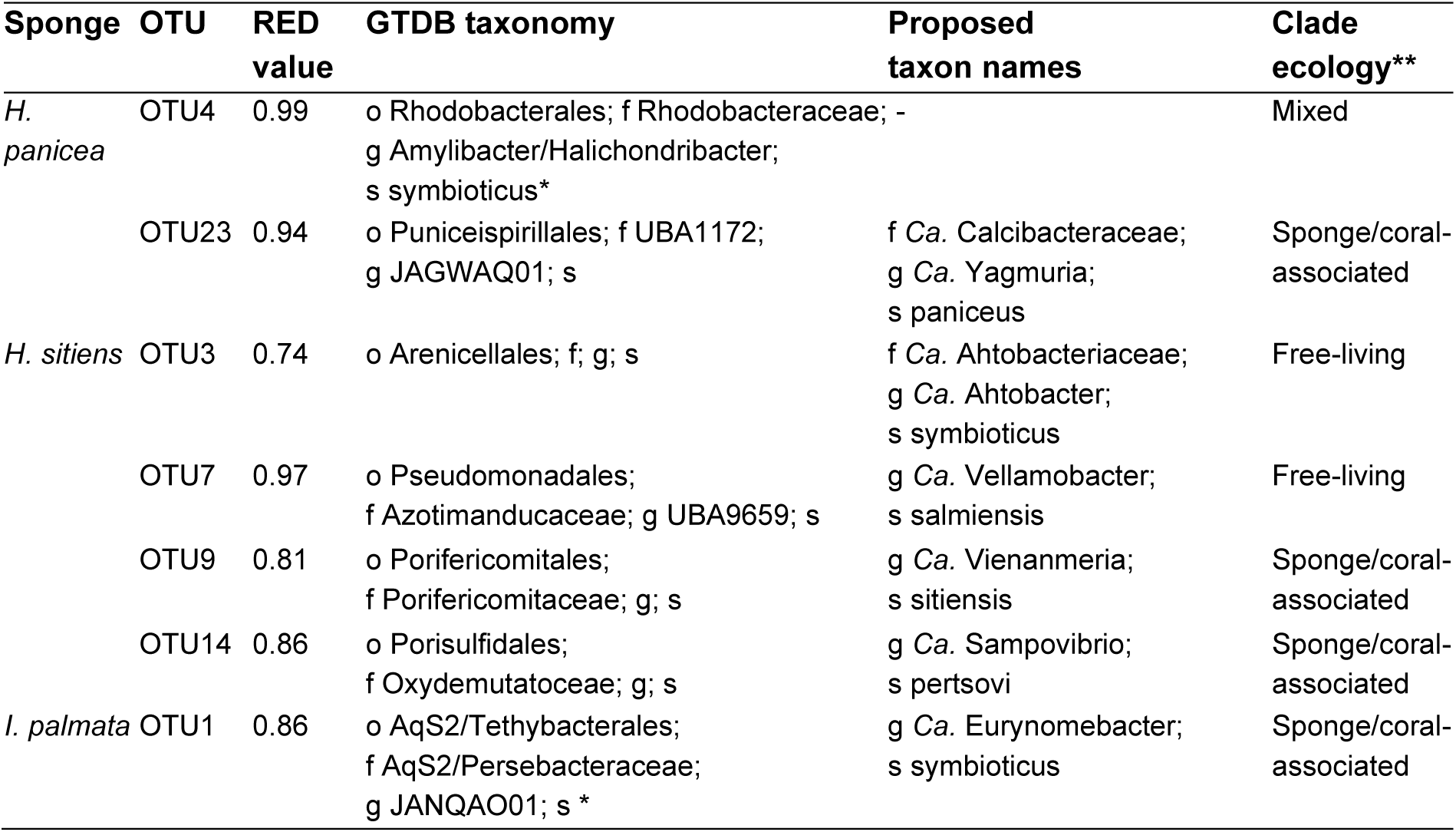
Classification of the identified bacterial sponge symbionts using GTDB and the proposed naming for novel taxa. *The GTDB taxonomy was extended with taxa proposed in [67] and [104]. **A bacterial clade was assigned to sponge/coral-associated if bacterial sequences were predominantly isolated from these animals; to free-living if sequences were predominantly obtained from marine sediments or water; to mixed if a fraction of sequences was obtained from sponges or corals.

The dominant HS-associated OTU3 MAG belonged to the order Arenicellales (Gammaproteobacteria) and represented the inaugural member of a novel family based on GTDB-derived taxonomy and a relatively low RED value (**Table 2**, **Figure 8**). Phylogenetic reconstructions based on a full-length 16S sequence revealed two potential members of this family isolated from marine sediments (**Supplementary Figure 7**). We propose a family name *Ca.* Ahtobacteriaceae, with the species type *Ca.* Ahtobacter symbioticus for the OTU3 bacterium, named after the sea god Ahto from Karelo-Finnish folklore. The species epithet “symbioticus” is related to the OTU3 lifestyle in the HS sponge.

**Figure 8.**
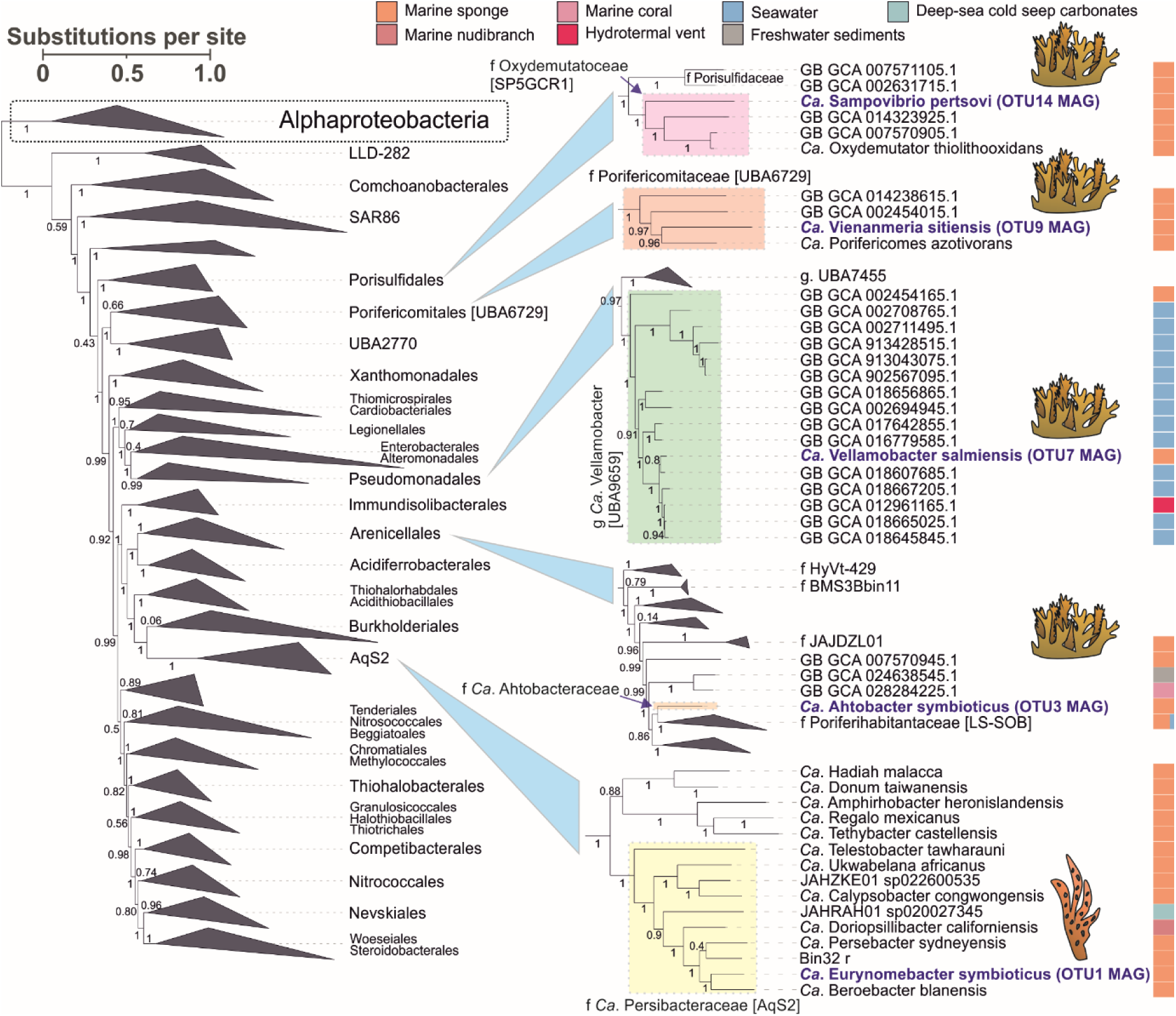
Maximum-likelihood tree of Gammaproteobacteria. The Alphaproteobacteria clade is shown as an outgroup. Local support values obtained with the Shimodaira-Hasegawa test are indicated for nodes (1000 resamples). Sponge-associated MAGs obtained in this study are highlighted in blue. Host sponges are indicated with cartoon icons. The isolation source for MAGs is color-coded.

The HS-associated OTU7 MAG was assigned to the genus UBA9659 (*Ca.* Azotimanducaceae/HTCC2089, Gammaproteobacteria), comprising primarily free-living bacteria from seawater (**Figure 8**). Correspondingly, sequences related to the OTU7 full-length 16S were isolated from seawater and freshwater environments (**Supplementary Figure 8**). We propose a replacement name *Ca.* Vellamobacter for the UBA9659 genus and species name *Ca.* Vellamobacter salmiensis for OTU7 bacteria. The genus name is derived from “Vellamo”, a goddess of water in Karelo-Finnish folklore. The species name indicates a place (the Great Salma strait of the White Sea) where samples were collected (**Table 2**).

The HS-associated OTU9 MAG was classified as a representative of a new genus within the Porifericomitaceae family, order *Ca.* Porifericomitales/UBA6729 (Gammaproteobacteria), a group known for its association with sponges [103] (**Figure 8**). Correspondingly, OTU9 full-length 16S belonged to *Ca.* Porifericomitales and was most close to a sequence isolated from *Hymeniacidon heliophila* sponge (95% identity) (**Supplementary Figure 9**). For OTU9, we propose a new genus name *Ca.* Vienanmeria and the species name *Ca.* Vienanmeria sitiensis. The genus name is derived from “Vienanmeri”, Karelian and Finnish name for the White Sea, while the species name reflects the host in which it was identified (**Table 2**).

The HS-associated OTU14 MAG was assigned to a novel sponge-associated family *Ca.* Oxydemutatoceae [103] within Porisulfidales [76] (Gammaproteobacteria) (**Figure 8**). Its full-length 16S sequence clusters with a 16S clone isolated from sponge *Ectyoplasia ferox*, which is basally positioned within *Ca.* Oxydemutatoceae (**Supplementary Figure 10**). We propose the name *Ca.* Sampovibrio pertsovi for OTU14. The genus was named after the “Sampo”, the magical object in the Karelo-Finnish epic Kalevala, believed to be a source of happiness and well-being. “Vibrio” reflects the morphology of this bacterium (**Figure 2F**). The species name honors Dr. N. A. Pertsov (1924-1987), the founder and director of WSBS MSU (**Table 2**).

IP-associated, OTU1 MAG was classified as a new member of the JANQAO01 genus belonging to the family of sponge and coral-associated bacteria recently described as *Ca.* Persebacteraceae (*Ca.* Tethybacterales, Gammaproteobacteria) [104] (**Figure 8**). Consistent results were obtained using 16S phylogeny (**Supplementary Figure 11**). We propose a new genus name *Ca.* Eurynomebacter as a replacement name for the JANQAO01 genus and the name *Ca.* Eurynomebacter symbioticus for the OTU1 bacteria specifically, following the naming logic proposed by Taylor et al. for Tethybacterales [104]. Eurynome is one of the Oceanids and a species epithet “symbioticus” reflects the ecological niche of this bacterium (**Table 2**).

Four out of seven studied bacteria belonged to clades of sponge/coral-associated bacteria, suggesting a symbiotic lifestyle and linked genomic adaptations. The other three belonged to mixed (*Ca.* H. symbioticus OTU4) or free-living (*Ca.* A. symbioticus OTU3 and *Ca.* V. salmiensis OTU7) groups of bacteria, presumably indicating evolutionary recent (or facultative) symbiosis with sponges.

## Discussion

In the present study, we deciphered the microbiomes of three species of Arctic LMA sponges. We identified a spectrum of increasingly complex associated bacterial communities consisting of one (IP), two (HP), and four (HS) species of symbiotic bacteria. We revealed that dominant symbionts were stably associated with sponge species over a 6-year period and across various geographical sites, while the minor symbionts may be site specific (for HP). Identified bacteria belong to two novel families, one novel genus, and three novel species of SABs from Alpha- and Gammaproteobacteria phyla.

Although SAMs demonstrated high stability over a long period of observation, samples from 2018 were compromised by an increased abundance of Alteromonadales and Vibrionales OTUs (**Supplementary Note 5**). Similar changes in microbiome composition, e.g., an increase in Alteromonadales and a decrease of SAB OTUs, were observed for the Antarctic sponge *I. kerguelenensis* after mechanical injury [70] and for HP after treatment with antibiotics [69]. Notably, mass mortality of sponges was detected in the summer of 2018 near WSBS MSU which was presumably linked to an abnormally high water temperature [28, 105]. Although cold-water sponges demonstrated some capacity to sustain stable microbiomes at elevated temperatures [65], our data indicate that acute thermal stress affects microbiome composition in a manner similar to tissue injuries or antibiotics. If so, a high relative abundance of Alteromonadales could serve as an early marker of sponge stress, preceding manifestation of visible phenotypic changes. We speculate that the elevated temperature negatively affects symbiotic bacteria which creates a niche for opportunistic bacteria such as Alteromonadales [106]. Alternatively, elevated temperature may stimulate the growth of Alteromonadales which causes infection of sponges and degradation of their native microbiomes. However, we did not observe proportionally increased abundance of Alteromonadales in the respective samples of sea water (**Supplementary Figure 1**). In the former scenario, the depletion of symbiotic bacteria precedes colonization and propagation of invading bacteria, while in the latter, invading bacteria actively hijack sponge holobionts and suppress bacterial symbionts. In both cases, the balance between symbiotic residential microbiome and opportunistic bacteria seems critical for sponge viability. Additionally, temperature can primarily affect the host causing its dysregulation and death [107]. The observation of symbiotic OTU in seawater and other sponges in 2018 may indicate a release of viable bacteria from decaying sponges. It should be noted that the sponge population at sampling sites seemingly recovered by the 2018 collapse. In 2022 (a year with a nearly normal temperature regime), studied sponge species were similarly abundant near WSBS MSU, and their microbiomes lacked signs of “invading” OTUs. Nevertheless, the observed perturbations highlight how fragile cold-water ecosystems are in the face of global warming.

Identified sponge symbionts are species-specific. Indeed, we observed their limited exchange between sympatric sponge species but without establishing new symbiotic interactions and symbiont proliferation in a new host. This suggests the existence of mechanism(s) allowing discrimination between self and non-self symbiotic bacteria even for closely related species of sponges (like, HP and HS). Discrimination and exclusion may be mediated by i) a resident symbiotic community (e.g., an active defense against invaders); ii) specific conditions in a host microenvironment (e.g., availability of specific metabolites or presence of viruses); iii) an interaction between a symbiont and a sponge (e.g., a specific recognition by the immune system or specific suppression by the immune system); or a combination of these factors. Natural products with antibacterial activities encoded by biosynthetic gene clusters (NP BGCs) are a widespread bacterial tool in competition for resources [108, 109]. Previously, diverse NP BGCs were reported in sponge symbiont MAGs, reaching 3.5 NP BGC per MAG [110, 111]. Symbiotic MAGs recovered in the current work were particularly poor in NP BGCs (<1 BGC/MAG), which makes the first hypothesis unlikely. However, this does not rule out the presence of yet unidentified NP BGCs that remain undetected by standard algorithms such as antiSMASH, nor does it exclude the possibility of alternative defense strategies. Although the exact conditions within the tissues of the different studied sponges are not known, transcriptomic, genomic, and ecological data indicate that they might be similar. Indeed, symbiotic MAGs recovered from different sponges showed similar metabolic patterns. As for the third scenario, mechanisms of interactions between bacterial symbionts and sponges are not studied yet. It was proposed that sponges may recognize specific bacteria by microbial-associated molecular patterns using their pattern recognition receptors, such as NLRs [58, 112, 113]. On the other hand, symbiont-encoded ELPs were considered as potential modulators of symbiont-host interactions [15, 95]. We found that symbiotic MAGs were significantly enriched in ELPs, and a substantial fraction of them (37%) carried a secretion signal. We hypothesize that such exported ELPs may be modulators of the host immunity system altering the response to general microbial-associated molecular patterns and licensing the presence of symbionts in sponge tissues or cells. Given the similar ecology of the three sympatric sponge species, which harbor relatively simple yet distinct SAMs, we propose them as a model system for further functional studies of sponge-symbiont interactions and specificity.

Earlier works on chemotaxonomic analysis of sponges used taurine content as a taxonomical marker [114, 115]. Particularly, both Halichondrida (includes HS and HP) and Poecilosclerida (includes IP) were characterized by high taurine content of >18% [83]. It has never been experimentally confirmed that sponges are responsible for taurine synthesis; however, transcriptomics and genomics of sponges indicated the presence of all necessary genes for this process. Several studies identified genes responsible for taurine import and processing in genomes of sponge symbiotic bacteria [14, 116–118]. Here, we further expand the association between sponge symbiosis and taurine metabolism capabilities by demonstrating the presence of complete pathways for taurine import and dissimilation in three taxonomically distinct species of dominant sponge symbionts. Metabolism of taurine is, however, widespread in free-living marine bacteria [85, 86, 119]. Particularly, 8% of bins retrieved from seawater metagenomes in the current study carried complete taurine metabolism pathways. Therefore, taurine catabolism capabilities are likely not an exclusive feature of symbiosis but may be critical for symbiotic species.

In addition to taurine dissimilation pathways, we detected two types of gene clusters presumably responsible for sulfoacetate uptake and dissimilation in genomes of three major symbionts. It is currently unclear how sulfoacetate is generated in sponge holobionts. Potentially, it can be excreted by sponges or produced by other members of the microbiome as a byproduct of taurine deamination [91, 120]. In the latter case, cross-feeding between bacterial species is expected. Additionally, sulfoacetate can be acquired from seawater where it is presumably synthesized alongside other organosulfonates by bacterioplankton [121]. Further research is needed to determine how widespread sulfoacetate metabolism is among different taxonomic groups of sponge symbionts. The sulfoacetate catabolic pathway may be an indicative, though not exclusive, feature of sponge symbionts, similar to taurine metabolism.

Analysis of sponge microbiomes featuring different numbers of bacterial symbionts allowed us to identify functional commonalities between microbiomes and a specialization within them at the species level. As such, all symbiotic microbiomes are likely to use carbohydrates as a primary source of energy via the oxidative pathways. All symbiotic genomes were enriched with ELPs; however, they had different types of ELPs even for symbionts of phylogenetically close HP and HS. Further, different complete biosynthetic pathways of amino acids and vitamins/cofactors were encoded in different symbionts, especially in the case of HS symbionts (most prominently, for polar amino acids and B vitamins). Finally, complete taurine and sulfoacetate catabolic pathways were found exclusively in genomes of dominant symbionts but not for minor symbionts. A specific example of specialization is the sulfur conversion of HS symbionts. While OTU3, OTU9, and OTU14 encoded various sulfur oxidation pathways, OTU7 encoded the assimilatory sulfur reduction pathway and may benefit from sulfate produced by other symbionts. Taken together, these observations are concordant with a microbiome model in which symbionts occupy minimally overlapping niches and cooperate rather than compete for resources [122, 123]. Further studies are needed to discover the roles of ELPs in the possible specialization of symbionts and identify potential cross-feeding between the species. Overall, our data suggest that bacterial symbionts may utilize taurine and/or sulfoacetate synthesized by sponges as a carbon and energy source while simultaneously providing them with amino acids and vitamins, thereby complementing host metabolism and contributing to the holobiont.

## Acknowledgements

Authors would like to acknowledge the personnel of the MSU White Sea Biological Station, particularly, Alexander Tzetlin and Tatyana Neretina for the opportunity to collect and process samples at WSBS, the WSBS SCUBA diving team for sample collection, and Dr. Boris Osadchenko for identification of sponge species. FISH was performed at the Core Centrum of the Institute of Developmental Biology RAS.

## Data availability

Raw sequencing data are publicly available in the NCBI Sequence Read Archive (SRA) under the Project Accession number PRJNA1211653. MAGs can be accessed via the BioSample accession numbers SAMN46784898-SAMN46784904. Scripts used in this study are available on GitHub (https://github.com/sutormin94/Sponge_metagenomes_2024), including the CORITES algorithm (https://github.com/sutormin94/CORITES).

## Author contributions

Study design - D.S., A.R. Samples collection and processing - A.R., V.F., D.S, A.T., V. M. Data analysis - A.R., V.M., M.R., D.M., D.S., M.E. Microscopy - Y.L., A.F. Cultivation experiments - A.T. Initial manuscript drafting - A.R., D.S. Manuscript review&editing – A.R., A.I., D.S. Funding acquisition - A.I., D.S. All authors have revised the manuscript.

