## Supplementary Notes & Figures for "Taxonomically different symbiotic communities of sympatric Arctic sponge species show functional similarity with specialization at species level"

### Extended Materials and Methods

#### *Bacterial fraction isolation and DNA extraction*

For isolation of a bacterial fraction from sponge tissue, a 1 $cm^3$ fragment of sponge tissue was fragmented by razor and forceps in 25 mL of marine water sterilized by filtering through a 0.22 µm filter (Sartorius). The obtained suspension was centrifuged at 200×g for 5 min to remove tissue fragments, spicules, and eukaryotic cells. The supernatant was collected and centrifuged again at 3500×g for 10 min. The resultant pellet was defined as a bacterial fraction and used for DNA isolation. To obtain marine water microbiome, 3 L of water was filtered using the 0.2 µm Sterivex filter (EMD Millipore). The fraction bound to the 0.2 µm membrane was defined as a bacterial fraction and was used for DNA isolation.

DNA was extracted from bacterial fractions of sponge samples using the Diatom DNA Prep kit (Galart Diagnosticum, Russia, catalogue number 100 D1024). For the bacterial fraction of marine water, settled in the 0.2 µm Sterivex filter, the filtering unit was opened, and the membrane was fragmented using the sterile razor. DNA was purified from the fragmented filter with the Diatom DNA prep kit. Then, to remove residual contaminants, DNA was extracted twice with phenol-chloroform (1:1 v/v) followed by chloroform extraction and precipitated in ethanol.

#### *High-throughput sequencing*

Preparation and sequencing of V3-V4 16S rRNA amplicon libraries were performed using standard degenerate primers fused with sequencing adapters (Illumina guide for 16S Metagenomic Sequencing Library Preparation, Part number 15,044,223 Rev. B and **Supplementary Table S2**) at Skoltech Genomics Core Facility using the 250 + 250 bp paired-end protocol with Illumina MiSeq. An exception is the samples from 2022 which were sequenced at Evrogen (Russia) using Illumina NovaSeq 6000 in the 250+250 bp paired-end mode. Shotgun libraries for samples from 2016 and 2018 were prepared using Illumina TruSeq kit and sequenced in the 150+150 bp paired-end mode using Illumina NextSeq 500 or Illumina HiSeq 4000 (Skoltech Genomics Core Facility). Additional shotgun libraries for 2016 samples and libraries for 2022 samples were prepared using the VAHTS Universal DNA Library Prep kit (Vazym) and sequenced in the 150+150 bp paired-end mode using the MGI DNBSEQ-G400 (at the BGI Sequencing Center).

Long-read sequencing libraries for samples from 2016 (*H. sitiens, I. palmata*) and 2022 (*H. panicea, H. sitiens, I. palmata*) were prepared using the NBD196 kit (ONT) and sequenced with the R9.4.1 flow cell (FLO-PRO002, ONT) on PromethION (at the BGI Sequencing Center). Additional long-read libraries for samples from 2022 were prepared using the Ligation Sequencing Kit 1D (SQK-LSK109, ONT) according to the standard protocol. Genomic DNA was subjected to End Repair and A tailing by NEBNext FFPE DNA Repair mix and NEBNext Ultra II End repair/dA-tailing module (NEB). Sequencing adaptors were ligated using NEBNext Ultra II DNA Ligation module (NEB) after AMPure XP (Beckman Coulter) purification. The final product was cleaned using an LFB buffer. MinION sequencing was performed using the R9.4 flow cell (FLO-MIN106D, ONT). Base calling was performed using Guppy v 6.4.6 in the high-accuracy mode.

*16S rRNA data analysis*

Raw forward reads were trimmed and filtered using Trimmomatic v. 0.39 (SE -phred 33 HEADCROP 17 ILLUMINACLIP:2:30:10 MINLEN:150) [1], and survived reads were processed with DADA2 pipeline v. 3.6.2 [2] giving amplicon sequence variants (ASVs). The ASVs were clustered using MMseqs2 v. 10-6d92c [3] at coverage > 0.95 and identity > 0.98, and representative sequences of clusters were further treated as operative taxonomic units (OTUs). OTUs were returned to DADA2, and taxonomy was assigned to OTUs using the SILVA collection [4]. Phyloseq package v. 1.30.0 [5] were used for further analyses.

*Assembly and annotation of shot-gun metagenomes*

Quality of Illumina or BGI reads was checked with FastQC [6]. Adapters were removed, and reads were filtered using Trimmomatic v. 0.39 (PE -phred 33 LEADING:3 TRAILING:3 ILLUMINACLIP:2:30:10 MINLEN:36) [1]. Survived read pairs and forward unpaired reads were assembled with SPAdes v.3.15.4 (with metaspades and k-mer length 55, 99, 127 options) for *de novo* assembly [7]. Adapters were removed from nanopore reads with Porechop v. 0.2.4 (--barcode_threshold 90) [8]. For long-read-only assembly, trimmed nanopore reads were assembled with Flye v. 2.8.1-b1676 (with --nano-raw or --nano-hq and --meta options) [9] and polished with Medaka v. 1.6.0 [10]. For hybrid assembly, trimmed short and long reads were assembled with SPAdes (with metaspades and k-mer length 33, 55, 99 options). The resultant assemblies were assessed using QUAST v. 5.1.0 [11].

*Metagenome assembly binning*

Metagenome assemblies were binned using MaxBin 2.0 v. 2.2.7 [12], CONCOCT v. 1.1.0 [13], MetaBAT 2 v. 2.12.1 [14], and binny v. 0.2 [15], all with default settings. Short reads and long reads were aligned to *de novo* assemblies using bwa mem v. 2.2.1 [16] and Minimap2 v. 2.24-r1122 [17], correspondingly. Initial bins were assessed using QUAST and CheckM2 v. 1.0.1 [18]. Ribosomal RNAs were predicted in bins with barrnap v. 0.9 [19] and taxonomy was assigned to bins using the GTDB-Tk v. 2.4.0 classify_wf function (GTDB database release 220) [20].

*CORe contigs ITerative Expansion and ScaffoLding refining algorithm (CORITES)*

First, a core set of contigs was built from contigs shared between the selected metagenomic bins. Next, the core set was iteratively expanded by other contigs derived from the binned assembly, using information from long-reads aligned to the assembly with minimap2. A contig was included in the core set if it was bridged with any of the core set contigs by the number of long reads exceeding the threshold value specified with the bridge_strength parameter. Usually, the iterative expansion module converged after 3-8 iterations without dramatic inflation of the initial core set. Inflation was observed only with low bridge_strength parameter values. The expanded core set was scaffolded with LongStitch v. 1.0.5 [21] using long reads. The scaffolded bin was additionally polished through alignment with the assembly graph generated by SPAdes. A pair of contigs from the bin was combined if their order can be unambiguously determined using the topology of the assembly graph. If such ordering exists, the two contigs were merged, and the gap between them was filled with the highest covered path in the assembly graph. The refinement algorithm was named CORITES (CORe contigs ITerative Expansion and ScaffoLding) after its iterative expansion module (https://github.com/sutormin94/CORITES). Final metagenome-assembled genomes (MAGs) were filtered to remove contigs shorter than 500 nucleotides and assessed using Quast and CheckM2.

*Phylogeny reconstruction based on full-length 16S gene sequences*

Full-length 16S rRNA sequences derived from MAGs of sponge-associated bacteria were searched using blastn against NCBI nt (released on 10/07/2023), NCBI 16S rRNA (released on 16/06/2023), and SILVA NR99 (v. 138.1) databases with default parameters if otherwise is not specified. Top 500, 100, and 200 hits were collected from the databases, respectively. For OTU1, several overrepresented species (*Bordetella holmesii*, *Bordetella parapertussis*, *Bordetella pertussis*, *Bordetella bronchiseptica*, *Bordetella hinzii*) were initially excluded from the search against the nt database. Representative sequences for these species were later manually added to a dataset. 16S sequences retrieved from different databases were combined and clustered using MMseqs2 with parameters "--min-seq-id 0.999 -c 0.999" to remove duplicated sequences. Representative sequences were aligned using MUSCLE (default parameters) in MEGA-X [22], and the edges of the alignment were trimmed to remove sparse regions. An initial maximum likelihood tree was constructed in MEGA-X based on the multiple alignments with 100 bootstrap iterations.

*Culturing of the sponge-associated bacteria and colony screening*

A sponge sample was washed 3 times with sterile seawater filtered 2 times through the 0.22 µm filter membrane (Millipore). A sponge fragment (~1 $cm^3$) free of other surface macroorganisms (such as seaweed or invertebrates) was moved to a Petri dish, containing 1 ml of sterile seawater, and fragmented with a sterile razor. The obtained sponge mass was processed with Dounce homogenizer, and the resultant homogenate was centrifuged for 5 min at 1000xg at 4°C to pellet the eukaryotic nuclei. The supernatant was collected, and a series of 5 consecutive 10-fold dilutions was made using sterile seawater. 100 µl aliquots of the initial supernatant and dilutions were plated onto poor solid medium (1.5% agar in sterile seawater), poor solid medium with yeast extract and taurine (0.04% yeast extract, 50 mM taurine, and 1.5% agar in sterile seawater, pH adjusted to 8.1 - 8.2), or rich solid medium with taurine (Bacto Marine Broth (DSMZ-Medium 514; Difco 2216), 50 mM taurine, and 1.5% agar in sterile seawater, pH adjusted to 8.1 - 8.2). Plates were incubated for 1-3 weeks at 4°C. Obtained colonies were reseeded on fresh plates with corresponding media and checked with PCR using symbiont-specific primers designed for the dominant sponge-associated MAGs (**Supplementary Table S2**).

Symbiont-specific primers were validated by PCR of metagenomic DNA extracted from the sponge samples. All primers gave PCR products with the expected length for corresponding sponge samples (data not shown).

*Fluorescent in situ hybridization (FISH)*

Probes specific to 16S sequences of identified SABs were designed using the Design Probes tool from DECIPHER with default hybridization conditions [23]. Probes with predicted high specificity (scores close to 0) were evaluated with SILVA TestProbe 3.0 [24] to exclude cross-reactive sequences matching other taxonomic groups; sequences prone to form hairpins were also excluded. EUB338I, EUB338II, and EUB338III probes were used as universal bacterial probes [25]. Cy3-labeled SAB-specific and Cy5-labeled bacterial universal probes were synthesized in Evrogen (Russia) (**Supplementary Table S2**).

Collected sponge individuals were placed in aquariums with circulating seawater for 10-12 hours at 4ºC to remove associated invertebrates and debris. Tissue samples (1 $cm^3$) were dissected from the osculum region, the middle part, and the base of a sponge body. The tissue samples were fixed in a 4% formaldehyde in filtered sea water for 3-6 hours at 10-15ºC. Then, the samples were washed in PBS, dehydrated in chilled 50% ethanol and stored at -20ºC in 50% ethanol until further processing.

For hybridization, tissue samples were gradiently rehydrated by PBS and then were dissected into 3x3 mm fragments at room temperature. Fragmented tissue was washed with 200 ul of wash solution (45% formamide pH 7.0 (Acros organics Cat# 327235000), 300 mM sodium chloride, 30 mM sodium citrate, 0.5% Triton X-100) in a 1:3 w/v ratio for 20 minutes. Hybridization was performed in 200 ul of hybridization solution (4 nM each FISH probe, 45% formamide pH 7.0, 300 mM sodium chloride, 30 mM sodium citrate, 50% dextran sulfate, 1% Triton X-100, 10 µg sheared salmon sperm DNA, 10 µg *Escherichia coli* RNase-free tRNA) for 10 h at 45ºC in a humidity chamber. An equimolar mixture of EUB338I, EUB338II, and EUB338III probes was used to stain all bacterial cells. After hybridization, samples were washed three times at RT for 20 min with 200 ul of hybridization solution without probes, followed by wash solution and PBS. DNA was stained with 200 ul 0.1 ug/ml Hoechst-33342 solution for 3 min and then washed with PBS. Stained samples were mounted with Prolong Gold antifade (Invitrogen) for confocal imaging and placed on confocal plates (Thermo Scientific). Imaging was performed in the Airyscan mode (Huff, 2016) using a Zeiss LSM 800 laser scanning confocal microscope equipped with a Plan-Apochromat 63x/1.4 Oil lens at the Center of N.K. Koltsov RAS. For fluorescent dye CY3, the excitation laser wavelength was 543 nm with filters BP 495-550 + LP570; for Cy5, the excitation laser wavelength was 633 nm with filters BP570-620 + LP645; for Hoechst-33342, the excitation laser wavelength was 405 nm with filters BP420-480 + BP420-480 + BP 495-550. Digital zoom was set to 1.8 (minimum recommended for Airyscan mode) and the pinhole was set to 200 nm. Additional capturing was performed in z-stack (number of layers 3), and Tile scan modes.

**Supplementary Note 1.** Refining of metagenomic bins with CORITES.

The quality of MAGs can significantly affect their functional and taxonomic analysis [26]. Multiple binners are available for the collection of contigs into metagenomic bins [27]. They, however, demonstrate different performances with different metagenomes and different types of sequencing data [28, 29]. To leverage this problem, pipelines that combine output from different binners were developed [30–32]. Two simple procedures are usually utilized to merge related bins: a) only contigs shared between related bins are kept; b) related bins are ranked by quality and the best bin is kept. Both methods have obvious limitations. Filtering of contigs reduces binning errors but at a price of incomplete genomes obtained (too strict merging). Ranking of bins results in high-quality bin selection but at a price of false assignment as no additional bin validation methods are typically included in the pipelines (too loose merging). Both too strict and too loose merging may dramatically affect the outcomes from a subsequent bin analysis based on comparative genomics such as the presence/absence of particular genes, metabolic pathways, etc. To address this problem, we developed the CORITES algorithm which aims to combine good properties of the above-mentioned bin merging strategies. In this algorithm, the initial core set of contigs is obtained by a strict merging which allows to reduce the chance of false contig assignment and bin contamination. Using the connectivity data (long reads or, potentially, Hi-C data) it then tries to expand the core set to increase the completeness of a MAG. The expansion procedure may result in contamination, especially in the case of a mixture of related genomes where connectivity data can erroneously link contigs originating from different genomes. For the sponge metagenomes, however, the CORITES algorithm demonstrated a good performance (especially, for medium-to-high abundant bacteria) and allowed us to increase the quality of initial bins. Particularly, the *Ca.* Halichondribacter symbioticus (OTU4) MAG obtained with CORITES and a recently published complete genome for this bacterium (NCBI OY365741.1) shared highly similar lists of pathways and other functional properties indicating the sufficient completeness of the derived MAG.

**Supplementary Note 2**. Additional genomes of *Ca*. H. symbioticus used in the study.

Using a BLAST search with the full-length *Ca*. H. symbioticus 16S sequence recovered from the refined OTU4 MAG, we identified two single-chromosome genomes (OY365738.1 and OY365741.1) in the NCBI nt database, which were generated as part of the Aquatic Symbiosis Genomics Project (https://www.aquaticsymbiosisgenomics.org/). Both genomes were obtained from HP and classified as uncultured *Amylibacter sp*. 16S rRNA sequences from these genomes showed 99.9% identity with OTU4 sequence, and the genomes themselves were clustered with OTU4 bins using GTDB-Tk (**Supplementary Figure 3**). To the compare the taxonomic position of the OTU4 MAG, we also used two previously published genomes, *Ca*. Halichondribacter symbioticus HS2 and *Ca*. Halichondribacter symbioticus Hp-f2 [33] (**Supplementary Table S6, S8**). We used the OY365741.1 genome for functional annotation and comparison with the OTU4 MAG.

**Supplementary Note 3.** Analysis of sponge transcriptomes used in the study.

To investigate the metabolic potential of sponges, we re-analysed publicly available transcriptomes Tr1 [34] and Tr2 [35] (for HP) and a transcriptome of *Isodictya sp.* [36]. After annotation and filtering, 35,943 proteins were predicted for *Isodictya sp.*, and 263,391 and 183,754 proteins were predicted for Tr1 and Tr2, respectively.

To determine whether any SAB genes were captured in the transcriptomes, we performed a BLAST search of predicted proteins from the OTU4 and OTU23 MAGs (HP) against the Tr1 and Tr2 transcriptomes and proteins predicted from the OTU1 MAG (IP) against the *Isodictya sp.* transcriptome. Using the identity and coverage thresholds of ≥95%, we detected the expression of multiple OTU4 and OTU23 SAB genes in the HP Tr1 transcriptome of HP. No SAB transcripts were found in the HP Tr2 and *Isodictya sp.* transcriptomes.

Throughout the analysis, we used the HP Tr1 transcriptome to detect the expression of bacterial genes and the HP Tr2 transcriptome to analyse the metabolic capabilities of the HP sponge, as it was not contaminated by bacterial transcripts. Similarly, we used *Isodictya sp.* transcriptome to predict metabolic capabilities of IP.

**Supplementary Note 4**. Analysis of metabolic pathways associated with taurine catabolism in MAGs of dominant SABs.

Dominant SAB MAGs (OTU1, OTU3, and OTU4) encoded alanine dehydrogenase (Ald), which may increase production of pyruvate required for the activity of Tpa, also generating NADH and ammonium. Ammonium could be next assimilated by glutamine synthetase (GlnA) for the synthesis of glutamine. Notably, expression of these genes (*ald*, *glnA*) was detected for the OTU4 SAB in the HP (Tr1) transcriptome. No genes responsible for the oxidation of sulfite to sulfate were detected in genomes of major SABs. OTU4 and OTU1 SAB MAGs carried *cysIJ* genes encoding a component of CysGIJ complex, which can reduce sulfite to sulfide as a part of the ASR pathway [37, 38]. Subsequently, generated sulfide may be used for biosynthesis of cysteine from serine via *cysE*/*cysK* pathway found in a complete genome of *Ca.* H. symbioticus or homocysteine via the *metZ* pathway (present in both OTU4 and OTU1 MAGs). OTU3 lacked *cysGIJ* genes but carried genes enabling sulfite transformation into trithionate (*dsrABL* genes). MAGs of minor SABs – OTU7 (HS) and OTU23 (HP) – also carried genes for inactivation of toxic sulfite - the ASR pathway in OTU7 and *soeABC* in OTU23.

**Supplementary Note 5**. Alteromonadales and Vibrionales were increased in SAMs in 2018.

In 2018, the relative abundances of Alteromonadales (Alteromonas OTU6 and Pseudoalteromonas OTU5) and Vibrionales (Aliivibrio OTU13 and Vibrio OTU16) were strongly increased in both sponge and water microbiomes (**Supplementary Figure 1B**). Interestingly, Vibrionales and Alteromonadales showed different abundance patterns: while Vibrionales were prevalent in seawater, Alteromonadales were more abundant in sponges. This can indicate preferential propagation of Vibrionales and Alteromonadales outside and inside sponges, respectively, or selective accumulation of Alteromonadales inside sponges by filtration. Propagation of Alteromonadales inside sponges is supported by the observation that some sponge samples collected in 2018 retained nearly intact microbiomes (similar to those observed in 2016 and 2022) and contained lower levels of “invading” OTUs despite being collected in close proximity to each other (**Supplementary Figure 1B**).

### Supplementary Figures

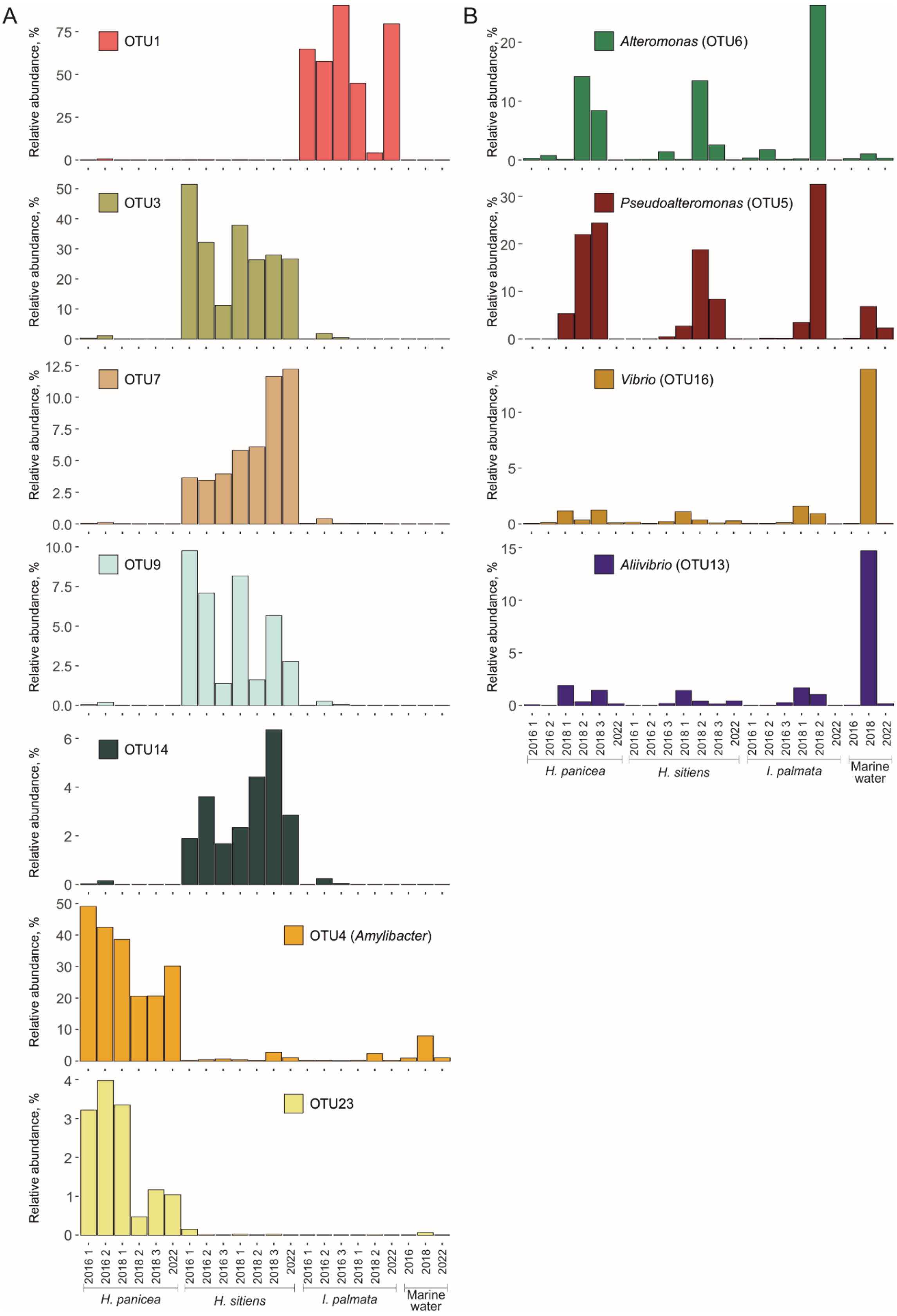


**Supplementary Figure 1.** Relative abundance of discussed OTUs in samples from the White Sea. (**A**) Relative abundances of sponge-associated OTUs. (**B**) Relative abundances of OTUs increased in samples from 2018.

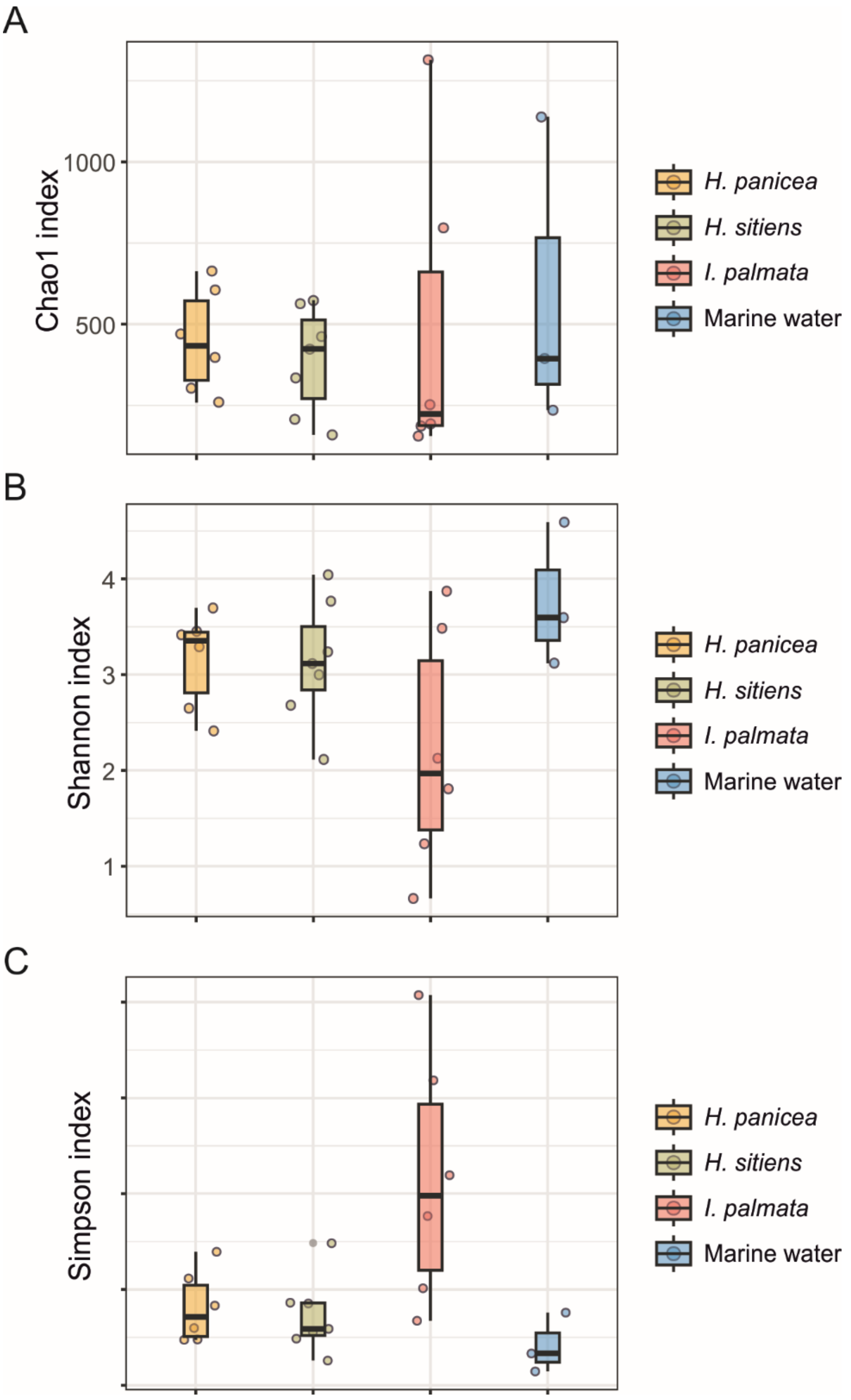


**Supplementary Figure 2**. Alpha-diversity indices for samples collected from different sponge species and the surrounding marine water from the White Sea. (**A**) Chao index. (**B**) Shannon index. (**C**) Simpson index. Individual samples are indicated with dots; median and quartiles are shown with box plots.

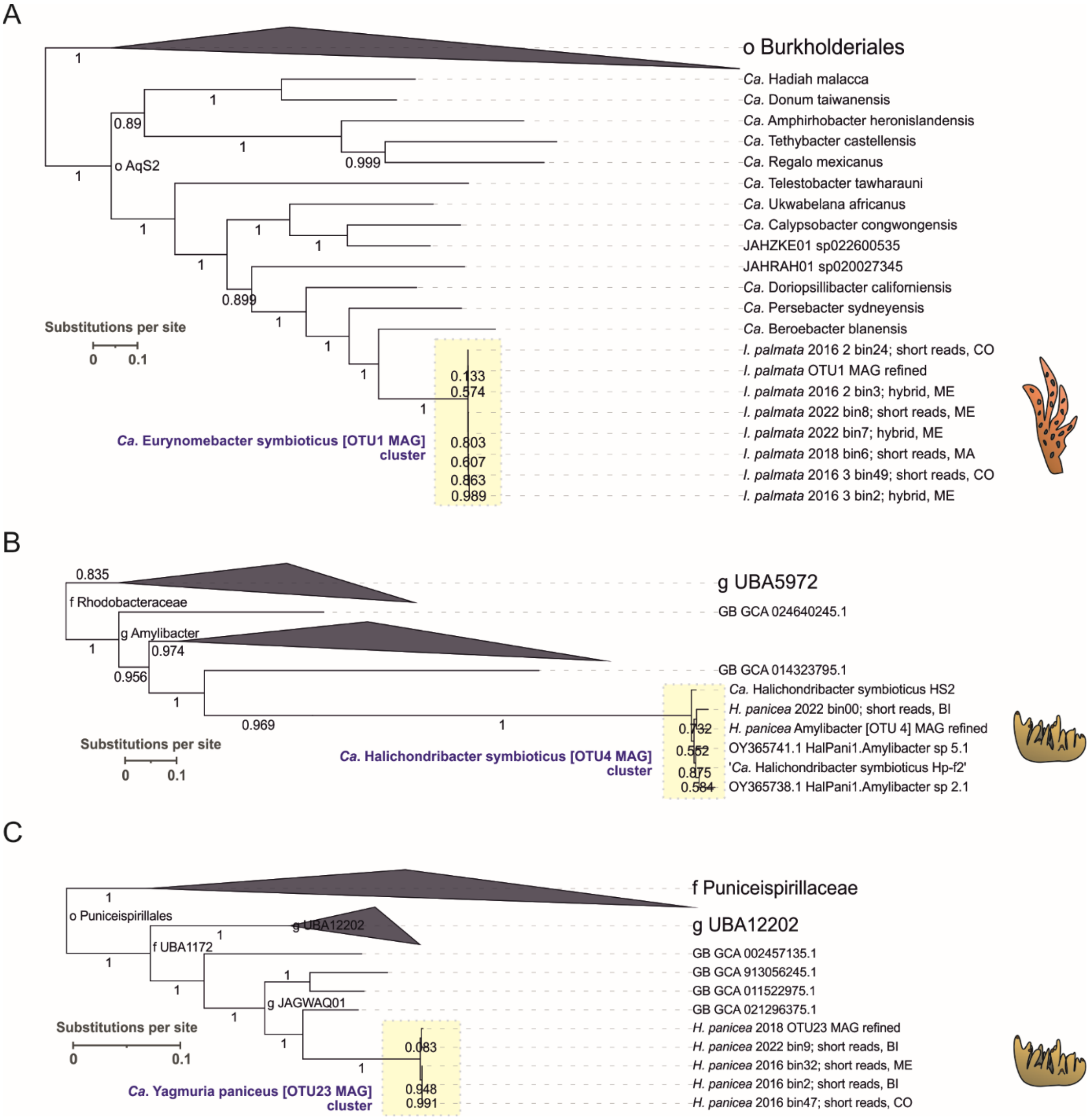


**Supplementary Figure 3**. Maximum-likelihood trees of SAB MAGs and bins obtained in this study with genomes available from GTDB constructed using GTDB-Tk. (**A**) OTU1 cluster, (**B**) OTU4 cluster (*Ca*. H. symbioticus), (**C**) OTU23 cluster, (**D**) OTU3 cluster, (**E**) OTU7 cluster, (**F**) OTU9 cluster, (**G**) OTU14 cluster. Local support values obtained with the Shimodaira-Hasegawa test are indicated for nodes (1000 resamples). Host sponges are indicated with cartoon icons. Clusters of SAB MAGs and bins obtained in the current study are highlighted with yellow boxes. For OTU4 (*Ca*. H. symbioticus), a *Ca*. Halichondribacter symbioticus Hp-f2 reference genome [33] was added, along with two single-chromosome genomes (OY365738.1 and OY365741.1) identified in the NCBI nt database. Notation of sponge-associated bins: sponge species from which a metagenome was obtained - year of sampling - bin ID; metagenome assembly strategy, binner was used. “Short reads” and “hybrid” indicate assembly from short read data and a hybrid assembly, respectively. CO - CONCOCT, BI - binny, ME - MetaBAT2, MA - MaxBin 2.0.

**Supplementary Figure 3**, continued.

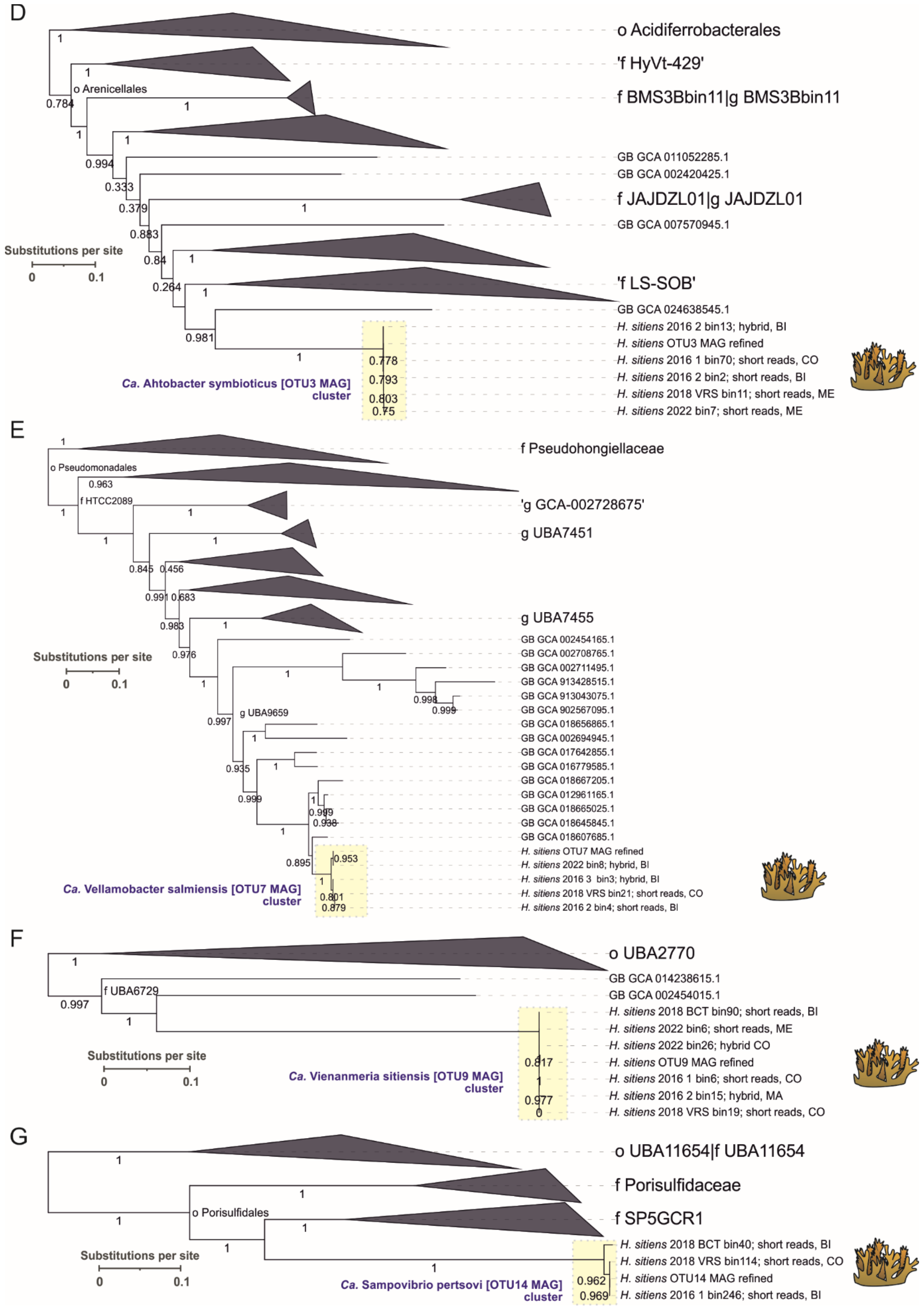

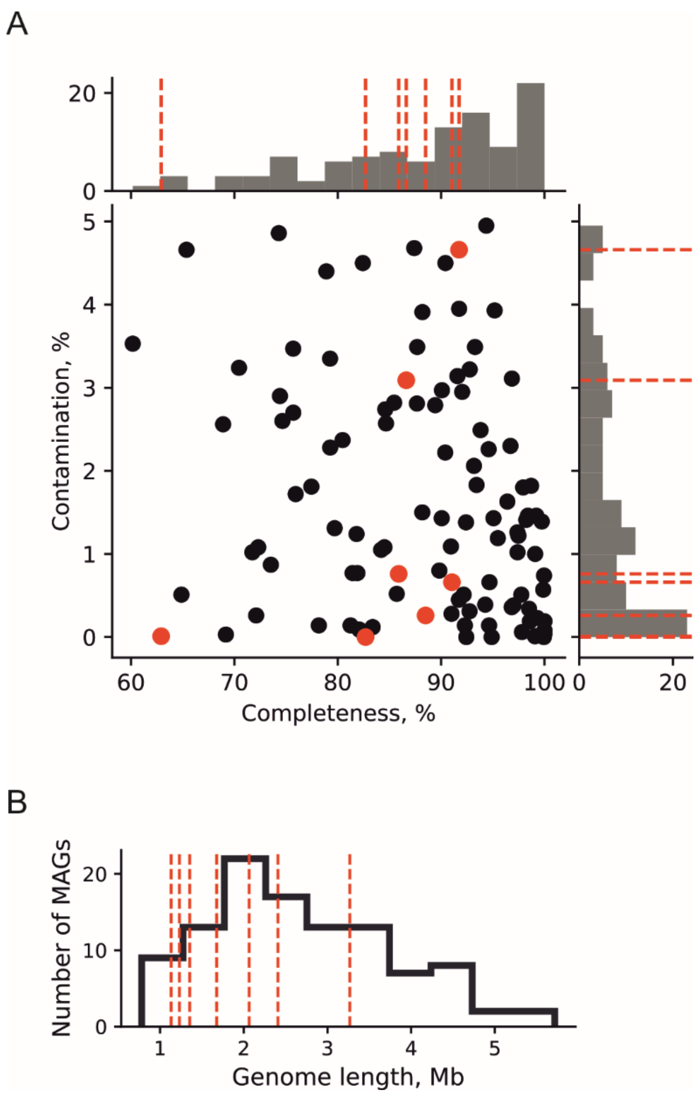


**Supplementary Figure 4**. Statistics of sponge-associated MAGs and other de-replicated metagenomic bins retrieved from seawater and sponge metagenomes. (**A**) Completeness and contamination of metagenomic bins and MAGs. Non-sponge-associated bins and sponge-associated MAGs are shown with black and red dots, respectively. (**B**) Distribution of total length of sponge-associated MAGs (shown with red vertical lines) and other metagenomic bins.

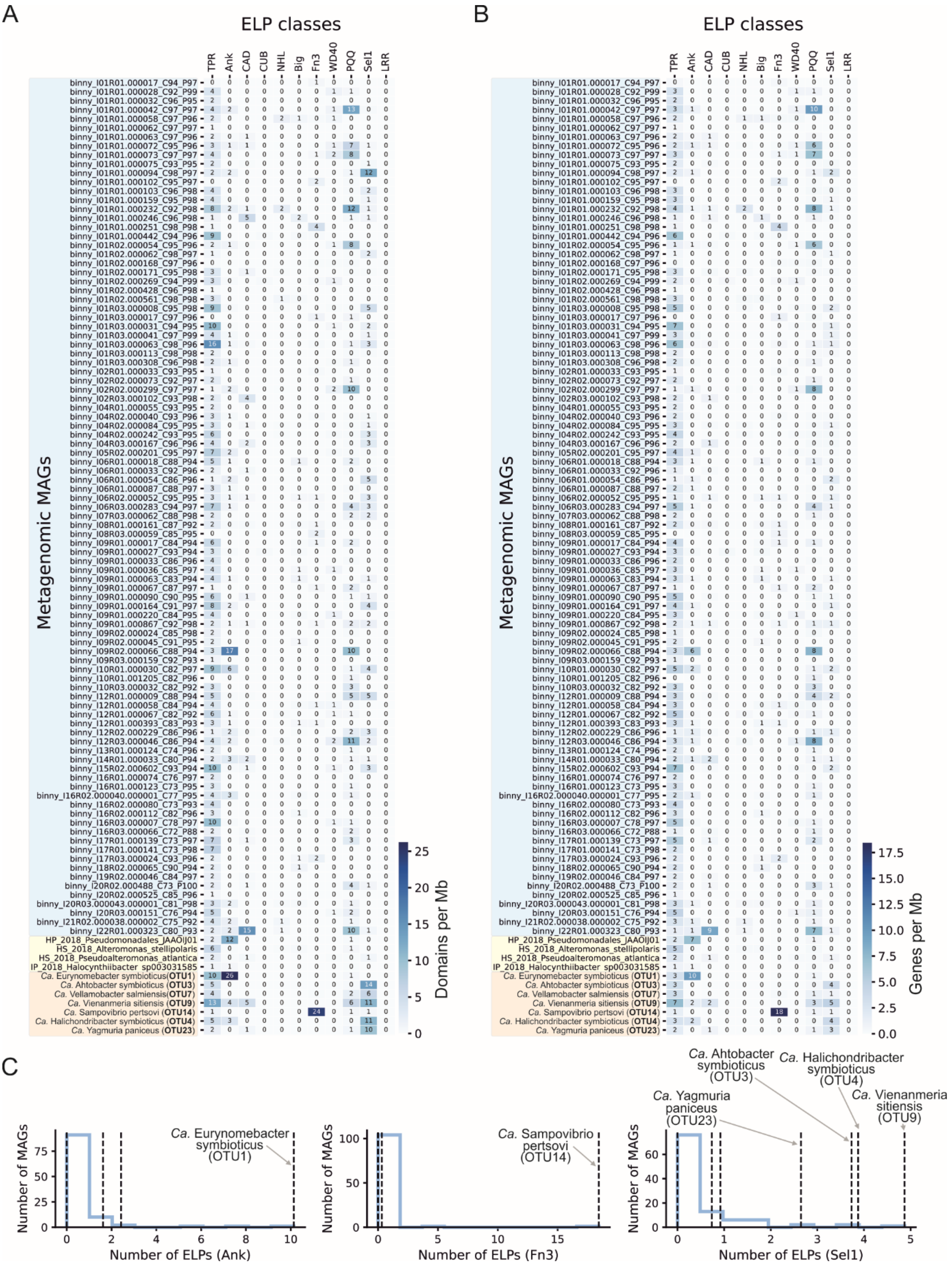


**Supplementary Figure 5**. Quantification of genes encoding eukaryotic-like domains in MAGs of sponge-associated bacteria. (**A**) A heatmap representing normalized frequencies of ELP domains (domain per Mb of a genome) in SAB MAGs (labelled with orange background), metagenomic bins of non-sponge-associated bacteria reconstructed from sponge metagenomes (yellow background), and bins from seawater metagenomes (blue background). (**B**) A heatmap representing normalized frequencies of ELP domain-containing genes (gene per Mb of a genome) in metagenomic bins (same as in panel A). (**C**) Distributions of normalized ELP-encoding gene frequencies in the analysed metagenomic bins and SAB MAGs. Frequencies of ELPs-encoding genes in SAB MAGs are indicated with vertical dashed lines. Data is shown for ELPs-encoding genes enriched in SAB MAGs.

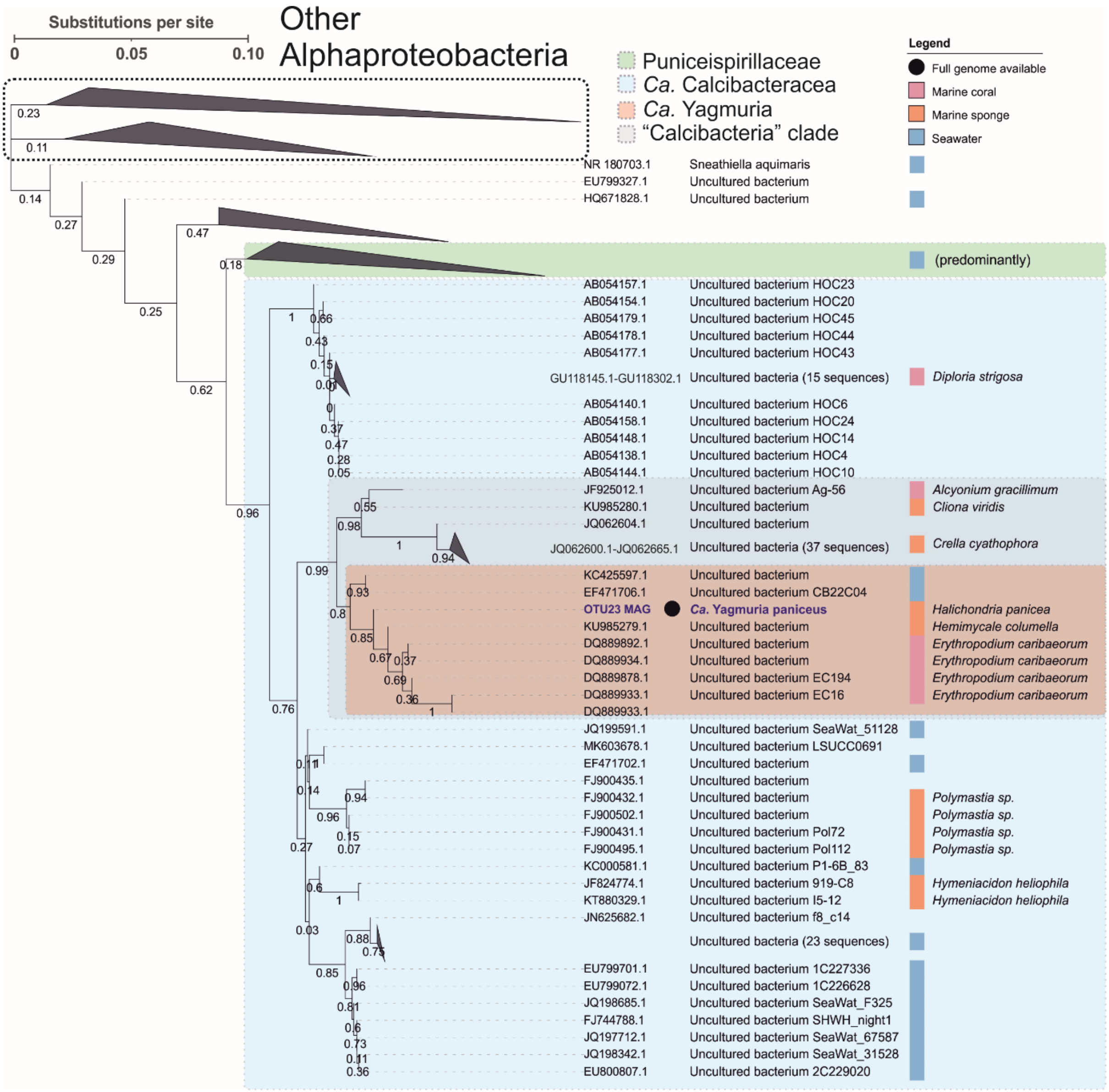


**Supplementary Figure 6**. Maximum-likelihood tree constructed for *OTU23* and related nearly complete 16S rRNA sequences from SILVA, rRNA NCBI, nt NCBI, and GTDB databases. The tree scale bar represents 10% sequence divergence. Bootstrap values are indicated on branches (100 replicates). For leaves, sequence accession number, bacterial species name, and species name of a host organism are indicated. The availability of a full genome sequence is indicated with a black dot. The biome type from where bacteria were isolated is indicated with a colour strip. The OTU23 sequence is highlighted in blue.

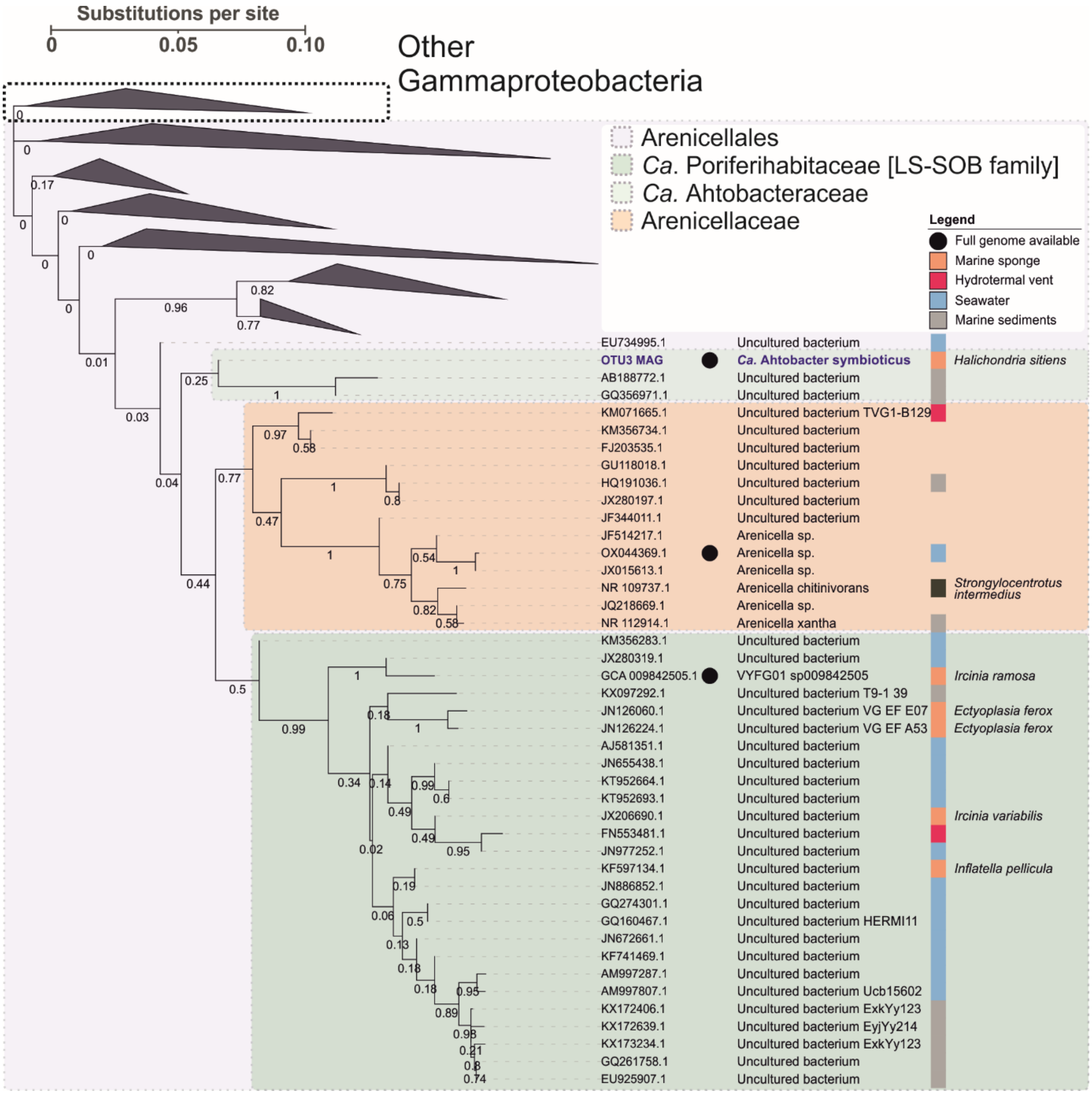


**Supplementary Figure 7**. Maximum-likelihood tree constructed for *OTU3* and related nearly complete 16S rRNA sequences from SILVA, rRNA NCBI, nt NCBI, and GTDB databases. The tree scale bar represents 10% sequence divergence. Bootstrap values are indicated on branches (100 replicates). For leaves, sequence accession number, bacterial species name, and species name of a host organism are indicated. The availability of a full genome sequence is indicated with a black dot. The biome type from where bacteria were isolated is indicated with a colour strip. The OTU3 sequence is highlighted in blue.

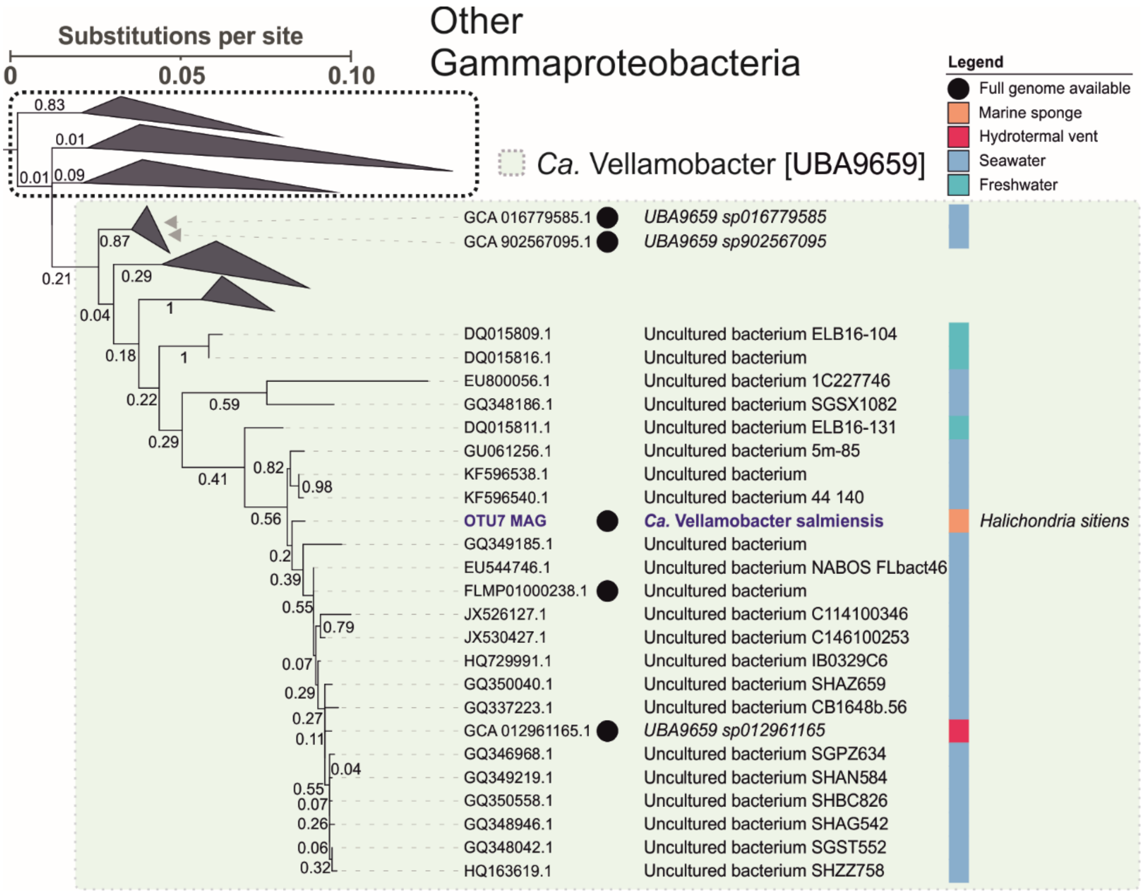


**Supplementary Figure 8**. Maximum-likelihood tree constructed for *OTU7* and related nearly complete 16S rRNA sequences from SILVA, rRNA NCBI, nt NCBI, and GTDB databases. The tree scale bar represents 10% sequence divergence. Bootstrap values are indicated on branches (100 replicates). For leaves, sequence accession number, bacterial species name, and species name of a host organism are indicated. The availability of a full genome sequence is indicated with a black dot. The biome type from where bacteria were isolated is indicated with a colour strip. The OTU7 sequence is highlighted in blue.

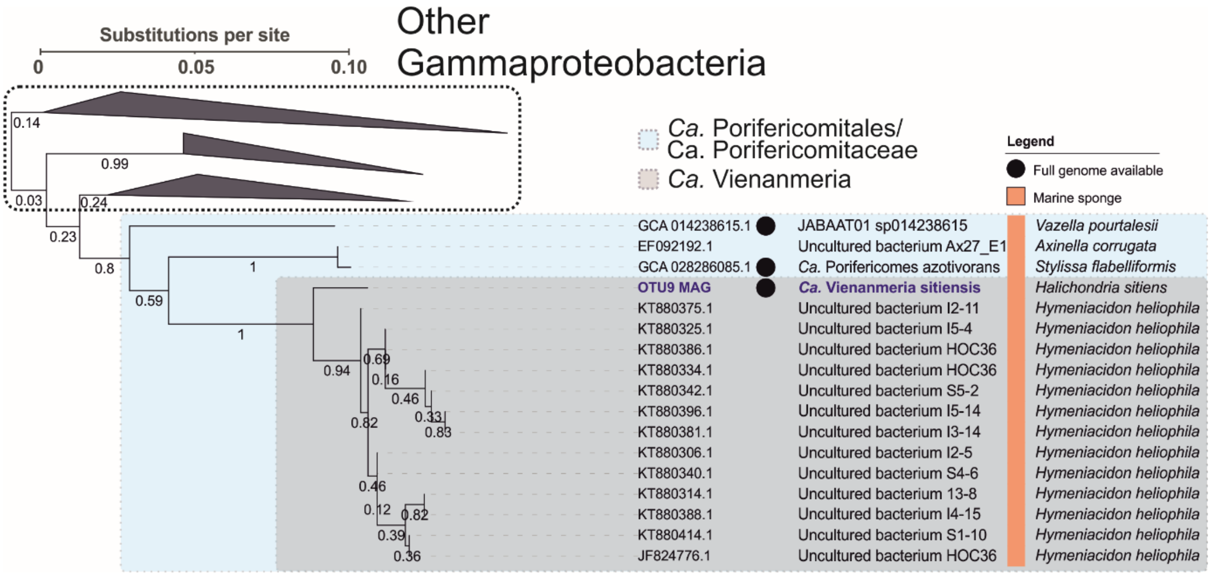


**Supplementary Figure 9**. Maximum-likelihood tree constructed for *OTU9* and related nearly complete 16S rRNA sequences from SILVA, rRNA NCBI, nt NCBI, and GTDB databases. The tree scale bar represents 10% sequence divergence. Bootstrap values are indicated on branches (100 replicates). For leaves, sequence accession number, bacterial species name, and species name of a host organism are indicated. The availability of a full genome sequence is indicated with a black dot. The biome type from where bacteria were isolated is indicated with a colour strip. The OTU9 sequence is highlighted in blue.

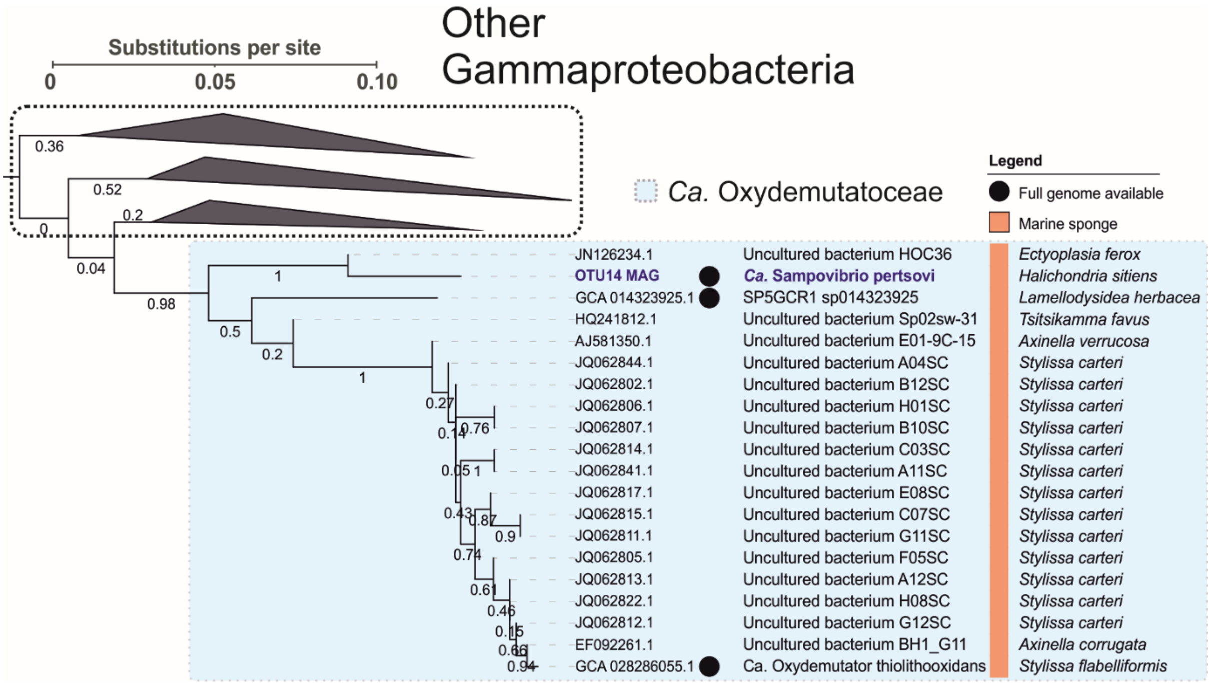


**Supplementary Figure 10**. Maximum-likelihood tree constructed for *OTU14* and related nearly complete 16S rRNA sequences from SILVA, rRNA NCBI, nt NCBI, and GTDB databases. The tree scale bar represents 10% sequence divergence. Bootstrap values are indicated on branches (100 replicates). For leaves, sequence accession number, bacterial species name, and species name of a host organism are indicated. The availability of a full genome sequence is indicated with a black dot. The biome type from where bacteria were isolated is indicated with a colour strip. The OTU14 sequence is highlighted in blue.

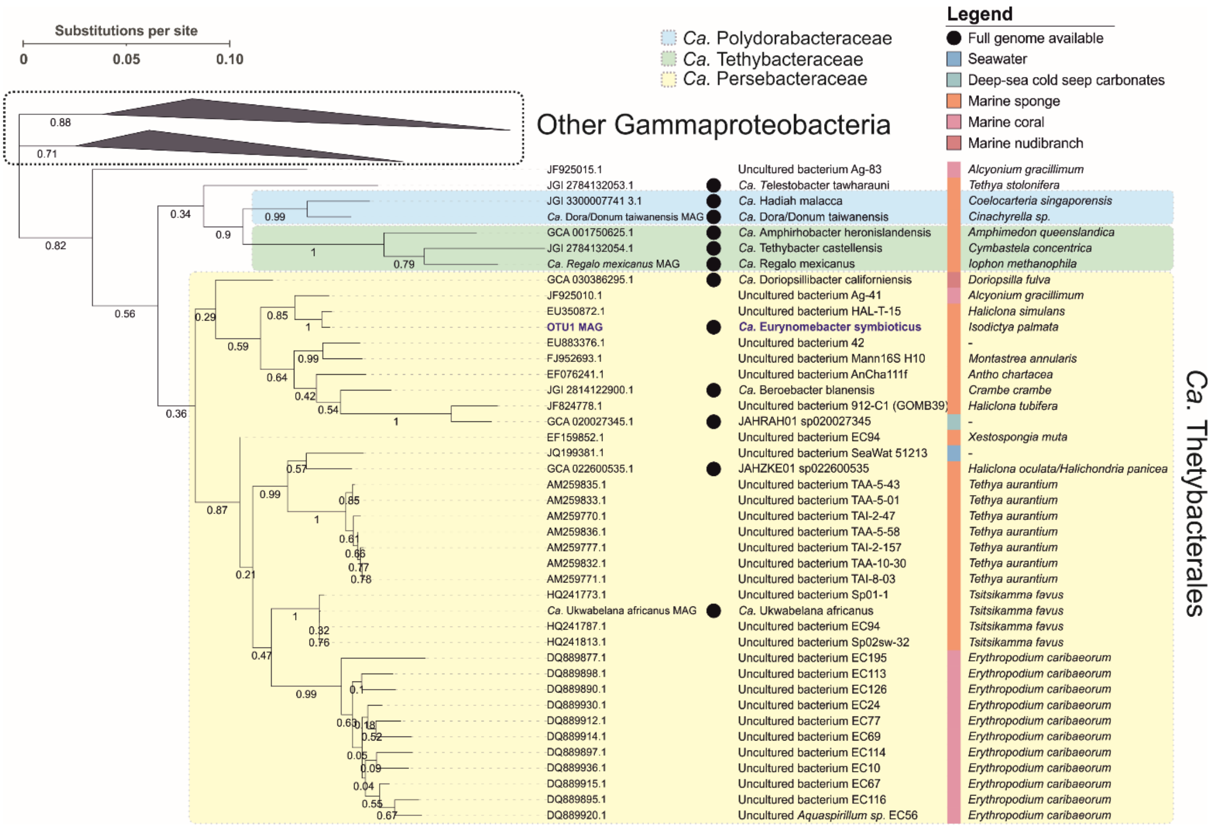


**Supplementary Figure 11**. Maximum-likelihood tree constructed for *OTU1* and related nearly complete 16S rRNA sequences from SILVA, rRNA NCBI, nt NCBI, and GTDB databases. The tree scale bar represents 10% sequence divergence. Bootstrap values are indicated on branches (100 replicates). For leaves, sequence accession number, bacterial species name, and species name of a host organism are indicated. The availability of a full genome sequence is indicated with a black dot. The biome type from where bacteria were isolated is indicated with a color strip. The OTU1 sequence is highlighted in blue.
